# The germline factor DDX4 contributes to the chemoresistance of small cell lung cancer cells

**DOI:** 10.1101/2022.04.22.489111

**Authors:** Christopher Noyes, Shunsuke Kitajima, Fengkai Li, Yusuke Suita, Saradha Miriyala, Shakson Isaac, Nagib Ahsan, Erik Knelson, Amir Vajdi, Tetsuo Tani, Tran C. Thai, Derek Xu, Junko Murai, Nikos Tapinos, Chiaki Takahashi, David A. Barbie, Mamiko Yajima

## Abstract

Human cancers often re-express germline factors, yet their mechanistic role in oncogenesis and cancer progression remains unknown. Here we demonstrate that DDX4, a germline factor and RNA helicase conserved in all multicellular organisms, contributes to epithelial mesenchyme transition (EMT)-like features and cisplatin resistance in small cell lung cancer (SCLC) cells. DDX4 depletion in H69AR and SHP77 cell lines decreased motility and resistance to cisplatin, whereas its overexpression increased these features. Proteomic analysis suggests that DDX4 upregulates metabolic protein expression related to DNA repair and immune/inflammatory response, suggesting its fundamental function may be in regulating cellular metabolism. Consistent with these trends in cell lines, DDX4 depletion compromised *in vivo* tumor development while its overexpression enhanced tumor growth even after cisplatin treatment in nude mice. Although the DDX4 expression level in somatic tumors is generally low compared to that in the germline, the relatively higher DDX4 expression in SCLC patients correlates with decreased survival and shows increased expression of EMT and cisplatin resistance markers. Taken together, we conclude that DDX4 influences the survival of SCLC patients by altering cellular metabolism in response to environmental cues such as drug treatments. This fundamental function of DDX4 as a germline factor might be applicable in other cancer types that express DDX4 and may serve as a key to combat specific tumors that are highly resistant to treatments.

**Highlights:**

- DDX4 contributes to cellular motility and drug resistance in SCLC cells.
- DDX4-overexpression globally alters the proteome and suppresses cytokine production.
- DDX4 promotes tumorigenesis and drug resistance *in vitro* and *in vivo*.
- DDX4 expression correlates with survival in SCLC patients and with immune/inflammatory response both in cell lines and patient samples.

## Introduction

Cancer cells can re-express germline factors typically restricted to gametes (Simpson et al., 2005). For example, when a malignant brain tumor is induced in *Drosophila* by inactivation of lethal (3) malignant brain tumor (l(3)mbt) protein, 25% of upregulated genes are germline factors. Further, inhibition of each germline factor (e.g. *vasa, piwi, aubergine*, or *nanos)* halted tumor growth, suggesting tumor dependency (Janic et al., 2010). The acquisition of germline characteristics in somatic cells may thus contribute to phenotypes characteristic of stem cells in general, and possibly tumorigenic cells. To date, few mechanistic studies have examined this biological phenomenon. Here we address this long-standing question in the field by focusing on the germline factor DDX4 in small cell lung cancer (SCLC).

DDX4 (*Drosophila* Vasa homolog) is one of the most conserved germline factors among all multicellular organisms. It often serves as a metric for germline determination because of its consistent expression across species, yet its function remains poorly understood. It is a member of the DEAD-box RNA helicase family (Raz, 2000; Linder, 2006; Gustafson and Wessel, 2010; Lasko, 2013), and found to be similar in sequence to eukaryotic initiation factor 4A (eIF4A) (Hay et al., 1988; Lasko and Ashuburner, 1988). It is considered to function as a regulator of mRNA translation in the germline (Hay et al., 1988; Lasko and Ashburner, 1988; Linder et al., 1989; Sengoku et al., 2006). Follow-up reports in *Drosophila* further suggest that Vasa associates with eIF5B, an essential translation initiation factor required for ribosomal subunit joining (Carrera et al. 2000; Pestova et al. 2000) and is involved in mRNA translation of germline-specific mRNAs in oocytes or germline stem cells (Johnstone and Lasko 2004; Liu et al., 2010). Further, more recent reports suggest that Vasa contributes to piRNA biogenesis and transposon silencing in mouse testes (Lim et al., 2013; Wenda et al., 2017). Therefore, for the last several decades, DDX4/Vasa had been thought to function exclusively in the germline.

Recent reports from our group as well as others, however, suggest that DDX4/Vasa might also function outside of the germline such as in embryonic development (Yajima and Wessel, 2011; Schwager et al, 2014; Fernandez-Nicholas et al. *in press*), in tissue regeneration (Wagner et al., 2012; Yajima and Wessel, 2015; see also review Poon et al, 2016). Vasa expression in these somatic cells is tightly regulated and often transiently expressed during a specific step of development. These observations suggest that DDX4/Vasa may be expressed in the soma only when linked to specific biological events, such as embryogenesis and regeneration, both of which require active cell proliferation and differentiation. This tight regulation of DDX4/Vasa expression in the soma implies that its uncontrolled expression may be pathogenic. Indeed, recent reports suggest that DDX4 plays an essential role in several human cancers including ovarian cancer (Hashimoto et al., 2008; Kim et al., 2014), multiple myeloma, and leukemia (Schudrowitz et al., 2017). Further, it appears to contribute to phase separation when expressed in the cells (Nott et al., 2015; Hondele et al. 2019). Based on these observations, we hypothesize DDX4/Vasa controls protein synthesis with spatial and temporal control, contributing to the rapid development and better survivability of the cells. However, the physiological significance of DDX4 in oncogenesis and tumor progression, especially its clinical implications remain poorly described.

SCLC is characterized by high rates of metastasis and relapse after treatment, leading to a dismal survival rate. No targeted therapy is approved for SCLC at current, partly due to a lack of molecular targets to explain the strong neoplastic features such as high rates of metastasis and chemoresistance. In this study, we demonstrate that DDX4 controls the cellular molecular landscape by facilitating mRNA translation, contributing to challenging clinical characteristics of SCLC. These findings provide an initial understanding of how germline factors contribute to cancer neoplasticity, a remaining question in the field while identifying a potential novel therapeutic target in DDX4-expression cancers.

## Results and Discussion

### DDX4 is expressed and enriched on the mitotic apparatus of SCLC cells

We previously identified several cancer types expressing DDX4 transcripts through database searches using the Cancer Cell Line Encyclopedia (CCLE) (Barretina et al., 2012) and Cancer Genomics Program (CPG, http://www.cancergenomicsprogram.ca/about-cgp). DDX4-high cancers include hematopoietic and neuroendocrine cancers (Schudrowitz et al., 2017). SCLC is a neuroendocrine cancer and SCLC cell lines were found on the DDX4 transcript-positive list, whereas lung adenocarcinoma cell lines were not found in our search. To confirm this database search result, we first profiled DDX4 mRNA and protein expression in several SCLC and NSCLC cell lines (Fig. 1). In all SCLC cell lines, DDX4 and another germline marker PIWI-like2 (PIWIL2) were detected by RT-PCR (Fig. 1A), and confirmed by sequencing of the PCR product (data not shown). DDX4 protein was found enriched on the mitotic apparatus during M-phase in those cells (Fig. 1B-C, arrows). Especially in H69AR and SHP77 cells that share a mesenchymal morphology and are partially adherent, DDX4 is stained in the cytoplasm and on the mitotic apparatus. DDX4’s localization to the mitotic apparatus was previously seen in hematopoietic cancer cells lines (Schudrowitz et al., 2017) as well as sea urchin embryonic cells (Yajima and Wessel, 2011), suggesting DDX4’s possible conserved function among various cells and organisms. In the NSCLC cell lines tested, on the other hand, DDX4 was barely detected by immunofluorescence or immunoblot (Fig. 1C-D). These results suggest that DDX4 may be expressed preferentially in SCLC, especially in the mesenchymal subtype.

**Fig. 1.**
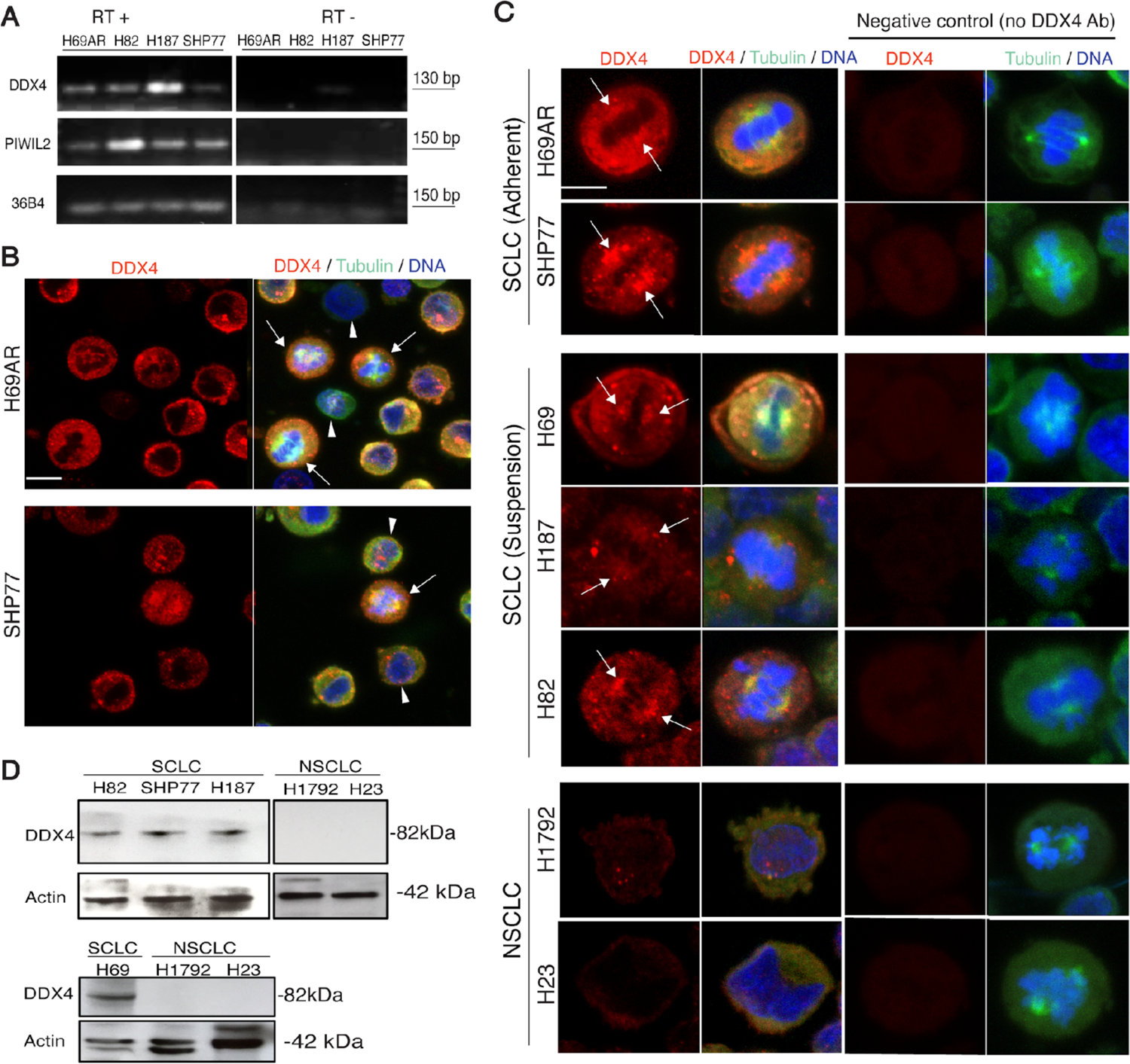
DDX4 is expressed in several SCLC cells. **(A)** RT-PCR results of four SCLC cell lines for three gene products (DDX4, PIWIL2, and 36B4), with negative controls (lacking RT enzyme) on the right, and positive tests on the left. (**B)** Immunofluorescence results and showing DDX4 expression in H69AR and SHP77 cells counter-stained with tubulin (green) and DNA (blue). Arrows indicate cells in M-phase. Scale bar = 5 μm. (**C)** DDX4 immunofluorescence results in five SCLC cells and two NSCLC cells counter-stained with tubulin and DNA. Arrows indicate the mitotic spindle. Images were taken at a time with the same experimental and laser condition for all cell lines. The signal level of DDX4 on the spindle or in the cytoplasm was calculated by *Image J* (n=10 each) and normalized to that of Tubulin. The average relative signal value equals or above 1 is indicated as +, the value above 1.2 is indicated as ++ on the right side of each cell image. Scale bar = 5 μm. (**D)** DDX4 immunoblot of various SCLC and NSCLC cell lines. DDX4 appears to be expressed in SCLCs yet not in NCSLC tested in this study. Actin was used as a loading standard.

### DDX4 depletion and overexpression change cell morphology in H69AR and SHP77 cells

To test the function of DDX4 in SCLC, we used some of those DDX4 expressing cells such as H69AR and SHP77 cell lines and constructed DDX4 overexpression (OE) cell lines that are driven by the EF1-α promoter along with controls that are introduced with NanoLuc (Figs. 2A and S1). DDX4-OE dramatically altered cell morphology with extended filopodia and a flattened shape with increased Cortactin expression which is important for lamellipodia formation (Fig. 2B-D).

**Fig. 2.**
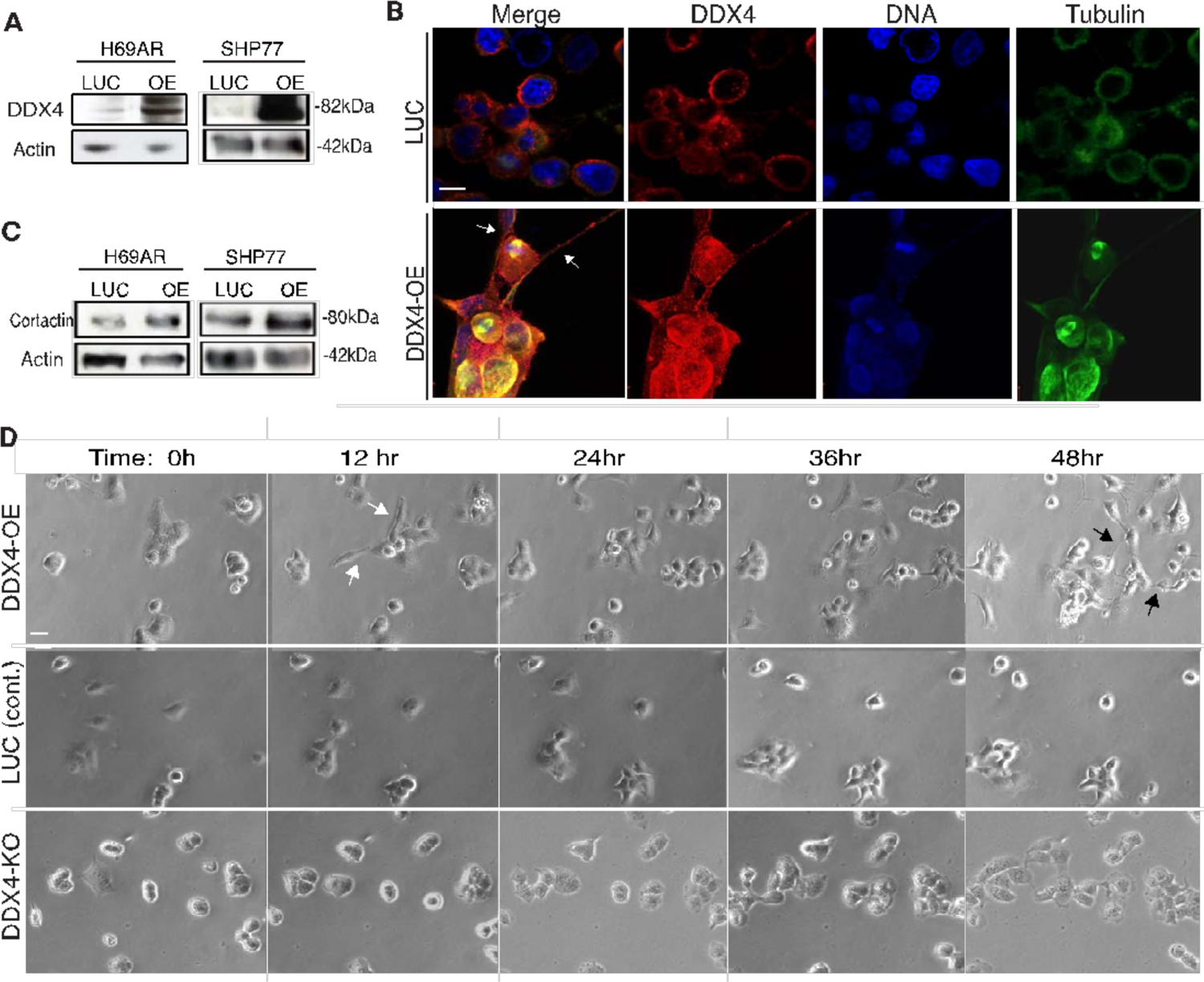
DDX4 overexpression (OE) induced the adhesive phenotype in H69AR and SHP77 cells. (A) Each cell line introduced with DDX4-OE transgenes showed an increased DDX4 protein level compared to the control (LUC). (B) Increased DDX4 protein level (red) was confirmed by immunofluorescence. DDX4-OE cell lines showed significantly different morphology characterized by flattened shape with extended filopodia. DNA, blue; Tubulin, green. Scale bar = 5 μm. (C) Cortactin protein level was upregulated by DDX4-OE both in H69AR and SHP77 cells. (D) Time-lapse images of DDX4-OE cell lines show random transformation into the mesenchymal state (white arrows) and increased cellular projections (black arrows), while DDX-KO cells stayed more rounded. LUC cells are shown as a control. Scale bar = 5 μm.

Similarly, we also constructed DDX4-knockout cell lines (Sg1-3) both in H69AR (Fig. S2) and SHP77 cells (Fig. S3) using CRISPR-mediated genome editing (Schudrowitz et al., 2017; see also Methods). The Sg1 and Sg3 lines showed slower cell proliferation compared to the control SC line (Fig. S2F), which is consistent with DDX4 depletion in multiple myeloma cells (Schudrowitz et al., 2017). CRISPR-mediated knockout (KO) occurs in a random manner, which may completely inhibit protein expression in some cells or may also result in truncated or mutated proteins that are functional in single-cell clones. To reduce this randomness, we further selected every five cells of the H69AR-Sg1 line that showed the most knockout efficiency (Fig S4A). Each of the isolated colonies was evaluated for DDX4-KO efficiency by genomic PCR and sequencing. As a result, the #2 colony group (DDX4-KO# 2) showed 100% mutation efficiency, and 81 % of those (n=21) caused a frameshift within or right after DDX4’s 3^rd^ exon (located between 128∼205 bp of the total 2073 bp of DDX4 ORF), significantly reducing DDX4’s expression in the population (Fig. S4B-C). Of note, we also attempted single-cell selection with 100% frameshift efficiency within the 3^rd^ exon of DDX4. However, DDX4-KO cells failed to proliferate after single-cell selection whereas control cells grew well under selection, suggesting that some minimal DDX4 expression is critical for cell survival. Although resultant KO efficiency varies among the three pooled clones of DDX4-KO cell lines (H69AR DDX4-Sg1∼3, SHP77 DDX4-Sg1∼3 and H69AR DDX4-KO# 2 and KO#1 lines), a similar phenotype was observed for all of them: DDX4-KO cells display a rounded cell shape with less adhesive feature compared to controls (Fig 2D; Movies S1-4). We, therefore, selected H69AR DDX4-KO# 2 and #1, and SHP77-Sg3 for further testing.

### DDX4 increases cell motility

To test whether DDX4 expression alters cellular motility, each group of H69AR cells was time-lapse imaged. The motility of each cell was automatically tracked by *FIJI*, which shows each cellular movement as an individual line: The longer line suggests more active cellular motility (Fig. 3A). As a result, DDX4-OE and –KO showed increased and decreased cellular motility, respectively (Fig. 3B-C). Similar trends were seen in SHP77 cells (Fig. S5). These results suggest that DDX4 increases cellular motility. Of note, we also attempted a conventional cell invasion assay using a transwell, yet DDX4-OE H69AR cells adhered to each other first and formed a tumor in the well, and which physically blocked them from penetrating the membrane under the condition we tested. Therefore, DDX4 appears to increase cell motility and cell-cell interaction.

**Fig. 3.**
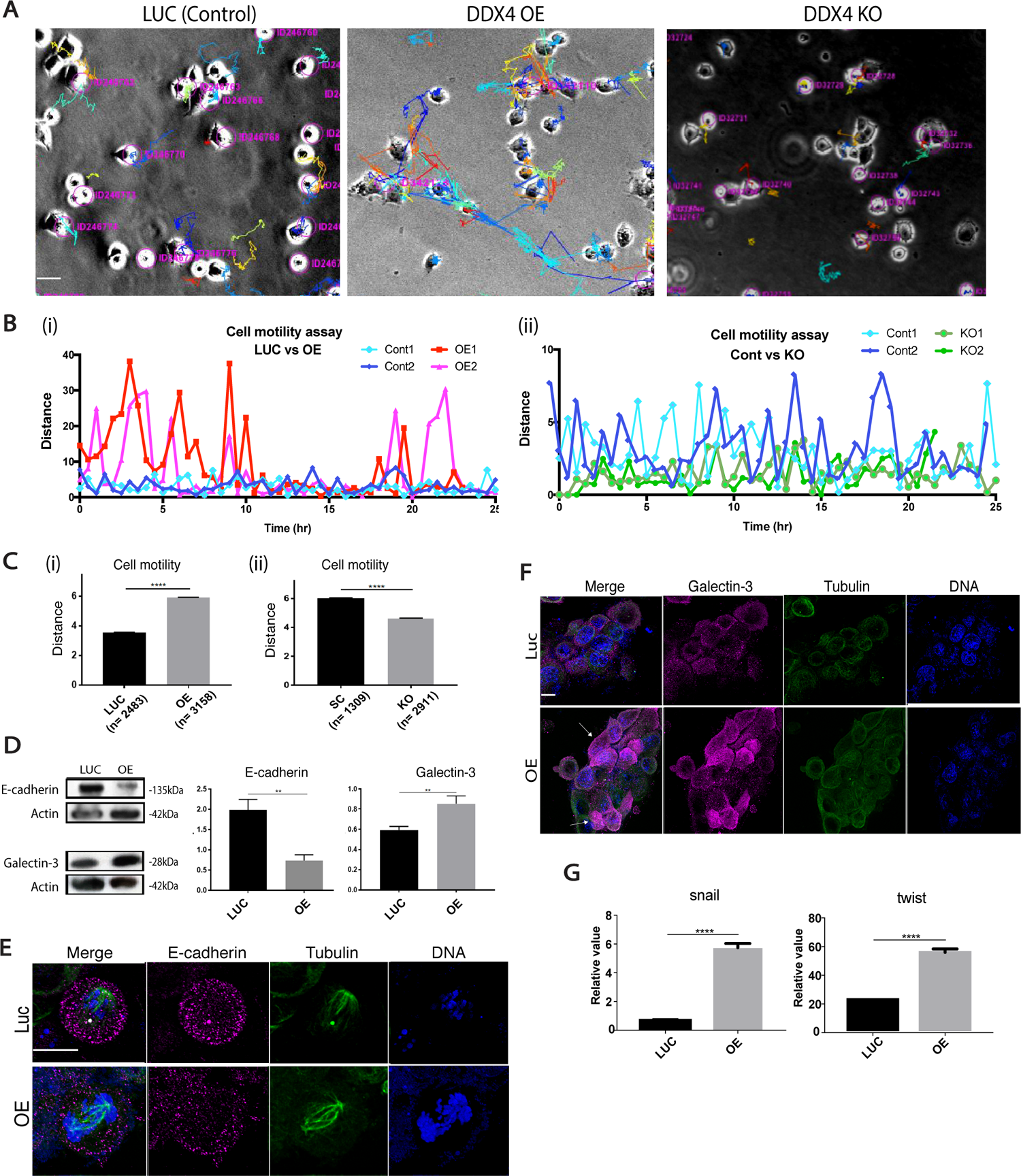
DDX4 expression increased cell motility and metastatic phenotypes. **(A-C)** Cell motility analysis of DDX4-OE and -KO lines in H69AR cells. Each cell line was timelapse-imaged every 30 minutes for ∼48 hours. Each colored line indicates a track of each cellular movement. The longer line suggests the larger movement (A). Cell motility analysis of a representative cell from each cell line. A distance of travel at each time point of time-lapse imaging was calculated and compared between DDX4-OE and LUC (left graph) and DDX4-KO and SC (right graph). DDX4-OE or –KO showed increased or decreased motility compared to each control, respectively (B). Travel distances of all cells in the field were determined by summating and averaging the distance value at each time point for each cell in the field. () indicates the total number of cells analyzed (C). The results show that DDX4-OE (i) and -KO (ii) cells traveled greater and lesser distances per 30 minutes, respectively. **(D)** Immunoblot results of E-cadherin and Galectin-3. The signal intensity of each protein was normalized by that of Actin to obtain the relative value shown in the graph. The graphs are the average of the four replicates. **(E-F)** Immunofluorescence of E-cadherin (E) and Galectin-3 (F) (magenta). Tubulin (green) and DNA (blue) were counterstained. The level of Galectin-3 level was highly heterogenic even within a population of DDX4-OE cells (arrows). Scale bars = 5 μm. **(G)** RT-qPCR results of *snail* and *twist*. The expression level of each gene was normalized by that of *h36B4*, a housekeeping gene. The graphs are the average of three replicates.

To evaluate whether DDX4 overexpression increases metastatic capacity, we tested expression levels of metastasis and angiogenesis markers in H69AR cells. In DDX4-OE cells, E-cadherin, a negative marker of metastasis, was downregulated, while positive markers such as Galectin-3 protein and the transcripts of *snail* and *twist* were all upregulated (Fig. 3D-G) (Nangia-Makker et al., 2018; Bure et al., 2019; Zhang et al., 2014). Taken together, these results suggest that DDX4 is critical for cell motility, and potentially contributes to metastasis of SCLC cells. Since DDX4/Vasa is considered to be involved in the collective migration of germ cells, it may similarly contribute to SCLC metastasis.

### DDX4 promotes chemoresistance

Platinum-based chemotherapy combined with Etoposide is the standard first-line treatment for SCLC, and most patients initially respond. Within a few months, however, SCLC often recurs, leading to devastating outcomes in most patients. To test DDX4’s contribution to cellular survivability, we examined the cell growth and viability of each H69AR cell line under treatment with several anti-cancer drugs including cisplatin, Hydroxide Urea (Hu), Camptothecin (CPT), and Etoposide (ETP). Cisplatin and Hu cause general DNA damage and inhibit DNA repair, respectively, and thus stop cell proliferation. CPT and ETP are both Topoisomerase inhibitors, inhibiting DNA unwinding and chromosome replication. We found that DDX4-OE decreased and DDX4-depletion increased sensitivity to cisplatin and Hu in a dose-dependent manner, while CPT and ETP showed no significant effect in any of the groups tested (Fig. 4A-B). Further, DDX4-OE decreased and DDX4-depletion increased phospho-γH2AX (DNA damage marker) slightly before cisplatin treatment, but more significantly, after cisplatin treatment (Fig 4C-E). Collectively, these results suggest that DDX4 increases resistance to cisplatin.

**Fig. 4.**
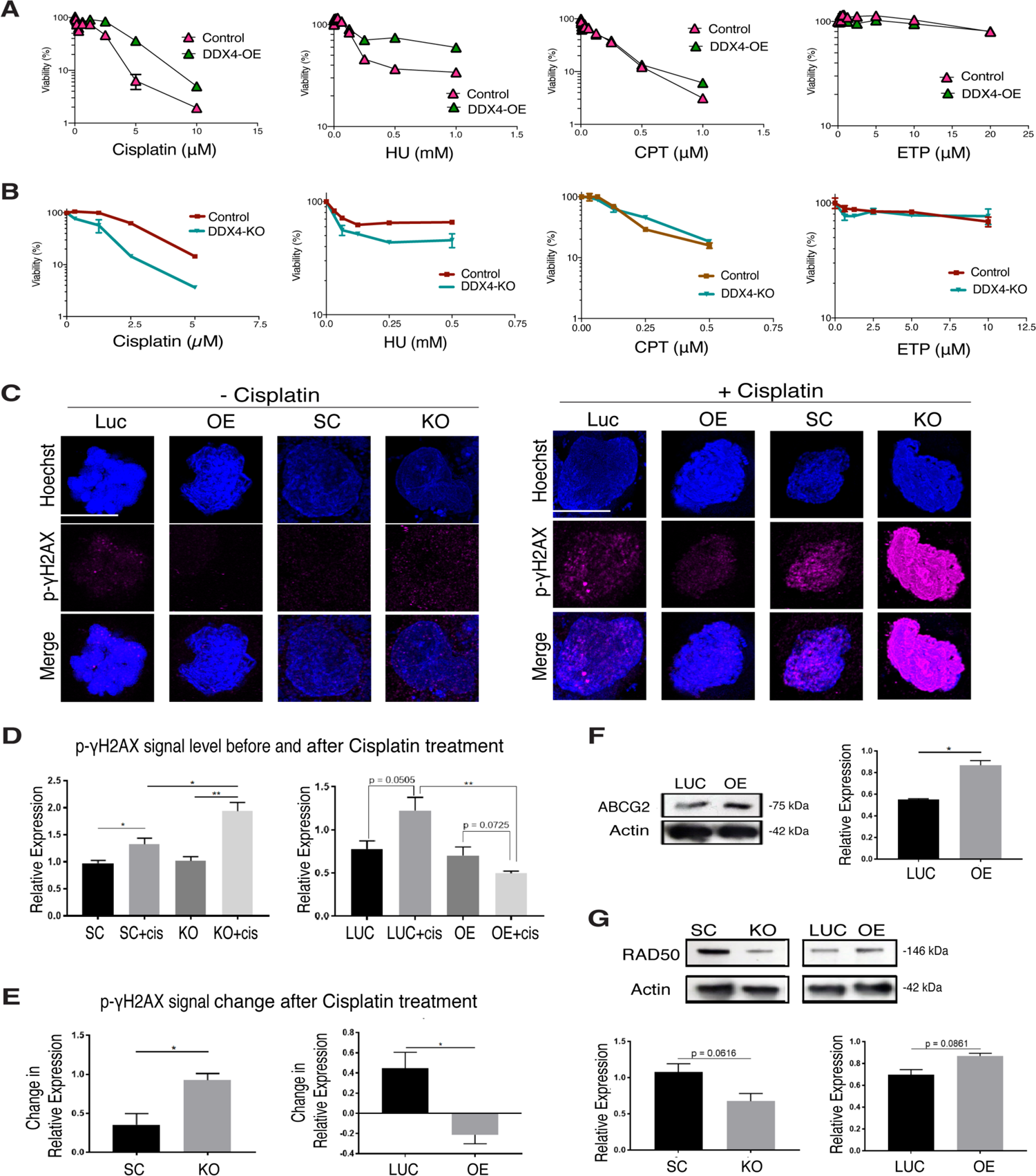
DDX4 promotes resistance to drug treatment and DNA damage. **(A-B)** Cell Viability (CTG) assay results of H69AR DDX4-OE (E) and -KO (F) cells. Each cell line was treated with various doses of drugs indicated in the X-axis for 3 days and the level of viability was measured. **(C-E)** H69AR cells were incubated with the final 10 μM of Cisplatin for 24 hours and then fixed. Phospho-γH2AX (magenta) levels were measured using immunofluorescence with levels being lower and higher in DDX4-OE and -KO, respectively. The graphs in Cʹ indicate the quantitative statistical analyses of the phospho-γH2AX signal before and after cisplatin treatment and the graphs in Cʹʹ indicate the level of signal increase after cisplatin treatment in each cell line. n= 8 ROIs. DNA, blue. Scale bar = 5 μm. (F-G) Immunoblot results of ABCG2 (E) and RAD50 (F). Both protein levels were slightly upregulated by DDX4-OE. The graphs indicate the average of 3∼5 replicates.

Further, we tested whether increased drug-resistance in DDX4-OE cells was due to increased DNA repair activity or clearance of these drugs by analyzing representative markers such as RAD50 and AGCG2: RAD50 is a DNA-repair molecule and AGCG2 is a member of the ABC-transporters responsible for pumping various foreign substances out of cells. A slight increase in both RAD50 and ABCG2 was found (Fig. 4F-G), implying that DDX4 may increase resistance to DNA damage by upregulating both the repair and pumping mechanisms, yet the underlying mechanisms warrant further investigation.

### Conserved function of DDX4 on the mitotic apparatus in somatic cells

To further support the above findings, we performed the rescue experiments using the wild type (WT) DDX4 and the C-terminus mutant (C3). This DDX4-C3 mutant contains three amino acid mutations in the C-terminus (Fig. 5A-B). This region is evolutionarily conserved yet distinct from other DEAD-box helicases (Fig. 5C) (Lasko, 2013), and was also recently reported to be critical for Vasa’s function in embryonic cells of the sea urchin (Fernandez-Nicholas et al., *in press*). We, therefore, hypothesize the C3 region is critical for DDX4’s activity in somatic cells.

**Fig. 5.**
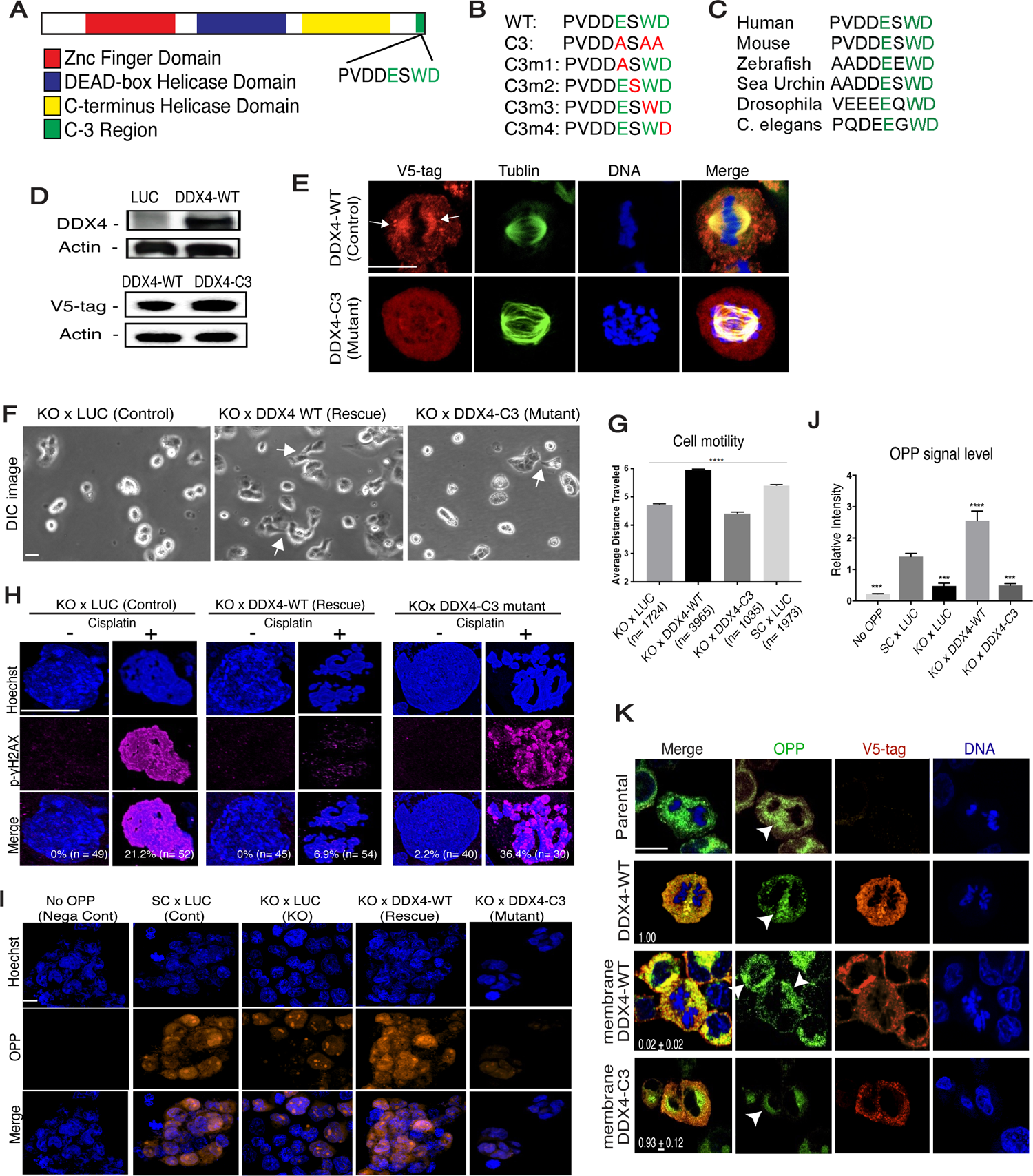
DDX4-C3 region is critical for restoring the DDX4-KO phenotypes in H69AR cells. DDX4-WT or DDX4-C3 mutant was introduced into the DDX4-KO cell line, tested for cell motility and resistance to DNA damage. **(A)** A schematic diagram showing the DDX4 molecular structure. The C3 region (green) is located at the end of ORF. (**B)** The amino acid sequence of the DDX4-C3 region is shown. The three amino acids colored in green (E, W, D) in the Wild Type (WT) were converted to Alanine (A) in the C3 mutant. One each of the three amino acids was converted to Alanine in a single-point mutant (C3m1∼4). (**C)** The sequences of the DDX4/Vasa-C3 region for various organisms are shown. The three amino acids colored green are highly conserved among organisms. **(D)** Immunoblot results of DDX4 and DDX4-C3. Upper panel, DDX4 antibody was used to detect DDX4. Lower panel, V5-tag antibody was used to detect DDX4-WT or DDX4-C3 to exclude the endogenous level of DDX4. Actin was used as a loading standard. **(E)** Immunofluorescence results of DDX4-WT and DDX4-C3 (red, detected by V5-tag antibody). Signal enrichment on the spindle (arrows) found in the DDX4-WT cell was little found in the DDX4-C3 cell. Tubulin, green; DNA, blue. Scale bar = 5 μm. **(H)** Each cell line was incubated with the final 10 μM of Cisplatin for 24 hours and then fixed for phospho-γH2AX (magenta) immunofluorescence. DNA, blue. % in the corner of each image indicates the proportion of the cells showing the positive p-γH2AX signal in each ROI. n indicates the total cell number analyzed in the 8 ROIs. To be noted, the signal level was in general higher in the control (LUC) and the C3 groups. **(F-G)** DDX4-WT rescued the phenotype with extended lamellipodia (F, arrows) and cell motility (G), while the DDX4-C3 mutant did not. Scale bar = 5 μm. **(I-J)** A level of mRNA translation was detected by OPP (orange). The signal intensity of OPP was normalized by that of DNA (blue) for each sample to obtain the relative signal level shown in graph Eʹ. The values are the average of the 8 ROIs through the three replicates. **(K)** Detection of translation (green, detected by OPP) in H69AR human cancer cell line introduced with DDX4 wild type (WT), membrane-targeted DDX4-WT, or membrane-targeted DDX4-C3 mutant (orange, detected by V5-tag antibody). The OPP signal was found enriched along the chromosomes or in the cytoplasm in the control (Parental and DDX4-WT) and membrane DDX4-C3 mutant groups but was found recruited to the membrane region in the membrane DDX4-WT group (arrowheads). Representative phenotypes are shown (n=10 ROIs). The number in the right bottom corner of each image indicates the average relative survivability of each cell line against the control line (DDX4-WT; set as 1) at two weeks after antibiotic selection (n=3). Scale bars =5 μm.

Both the DDX4-WT rescue line and the DDX4-C3 mutant line showed the same level of expression (Fig. 5D) yet its localization on the spindle was only found in the DDX4-WT cell line (Fig. 5E), suggesting that the C3 region may be critical for controlling DDX4’s subcellular localization. In those cell lines, we found that DDX4-WT rescued the mesenchymal morphology (Fig. 5F, arrows) as well as cell motility (Fig. 5G), while DDX4-C3 failed to rescue these features of H69AR cells. Consistently, DDX4-WT also rescued chemoresistance, while DDX4-C3 failed in H69AR cells (Fig. 5H).

Further, DDX4 is an RNA-helicase and may contribute to these phenotypes by rapidly changing protein expressions in the cells. To test this hypothesis, general mRNA translation of each cell line was analyzed by the signal level of O-propargyl-puromycin (OPP) that is incorporated into the active translational machinery and thus serves as a translation marker. DDX4-WT rescued translation, increasing it by 100% over the control, while DDX4-C3 showed a ∼70% decrease in translational activity similar to the negative controls (Fig. 5I-J). We also performed these rescue experiments using a DDX4-mutant with a single-point mutation in the C3-region (Fig. 4B, C3m1∼m4). Those single point mutants, however, equally rescued both mRNA translation and DNA-damage resistance capability to some extent (Fig. S6), suggesting that the three amino mutation is critical for DDX4’s activities in SCLC cell regulation.

Lastly, To test if DDX4’s localization on the spindle is important for its function in SCLC cells, we introduced membrane-targeted DDX4 (membrane-DDX4). Membrane-DDX4 caused ectopic translation at the membrane, and cells quickly died within one-two weeks after the construction of this cell line, even though endogenous DDX4 is present in those cells (Fig. 5K, OPP). The rest of the control cell lines including the membrane-targeted DDX4-C3 cells showed no aberrant translation nor defect in cell proliferation. These results support the idea that not the amount but the localization of DDX4 is critical for the survivability of SCLC cells.

Taken together, the fundamental role of DDX4 may be in controlling the subcellular location and timing of protein synthesis in the cell, contributing to neoplastic and metastatic features of SCLC cells. The evolutionary conserved C3-region appears to be the critical domain for this functionality of DDX4, suggesting a possible conserved mechanism of the DDX4 function across cancer and embryonic cells. Indeed, this C3 region is also reported to be important for Vasa’s function in piRNA biogenesis in the *Drosophila* germline (Dehghani and Lasko, 2015). Since PIWI, a major factor of piRNA biogenesis appears to be also expressed in SCLC cells (Fig. 1A), it will be important to test in the near future, if/how DDX4 contributes to piRNA and/or other small RNA biogenesis in somatic cancer cells.

### DDX4 changes metabolic protein expressions in SCLC cells

Since DDX4 appears to alter protein expression globally by affecting translation, this may allow cells to respond to external cues more effectively and increase cell survival. To test this hypothesis, we analyzed cytokine expression before and after cisplatin treatment in H69AR cells. We focused on cytokine expression because DDX4 expression dramatically changed the level of cell motility and cell-cell interaction in SCLC cells as shown in Fig. 3. As expected, the multiplex analysis showed that the cytokine expression profile was markedly different between the DDX4-OE and -depleted groups at baseline (Fig. 6A). After cisplatin treatment, changes in cytokine expression were found in both cell lines. In the DDX4-depleted group, the level of cytokine secretion was globally increased after treatment compared to the control group (SC) (Fig. 6B). RT-qPCR for cytokines and immune signaling components including STAT1 and CXCL10 also showed an upregulation in the DDX4-depleted group following cisplatin treatment, while the pro-survival factor IL-8 was downregulated after the treatment (Fig. 6C). On the contrary, upregulation of cytokine expression was globally repressed in the DDX4-OE group compared to the control (LUC) (Fig. 6B). These results further support the contention that DDX4 expression makes cells insensitive to cisplatin treatment. This will also likely help cells evade immune surveillance and survive during chemotherapy (Edwardson et al., 2017; Ni and Lu, 2018; Sen et al., 2019).

**Fig. 6.**
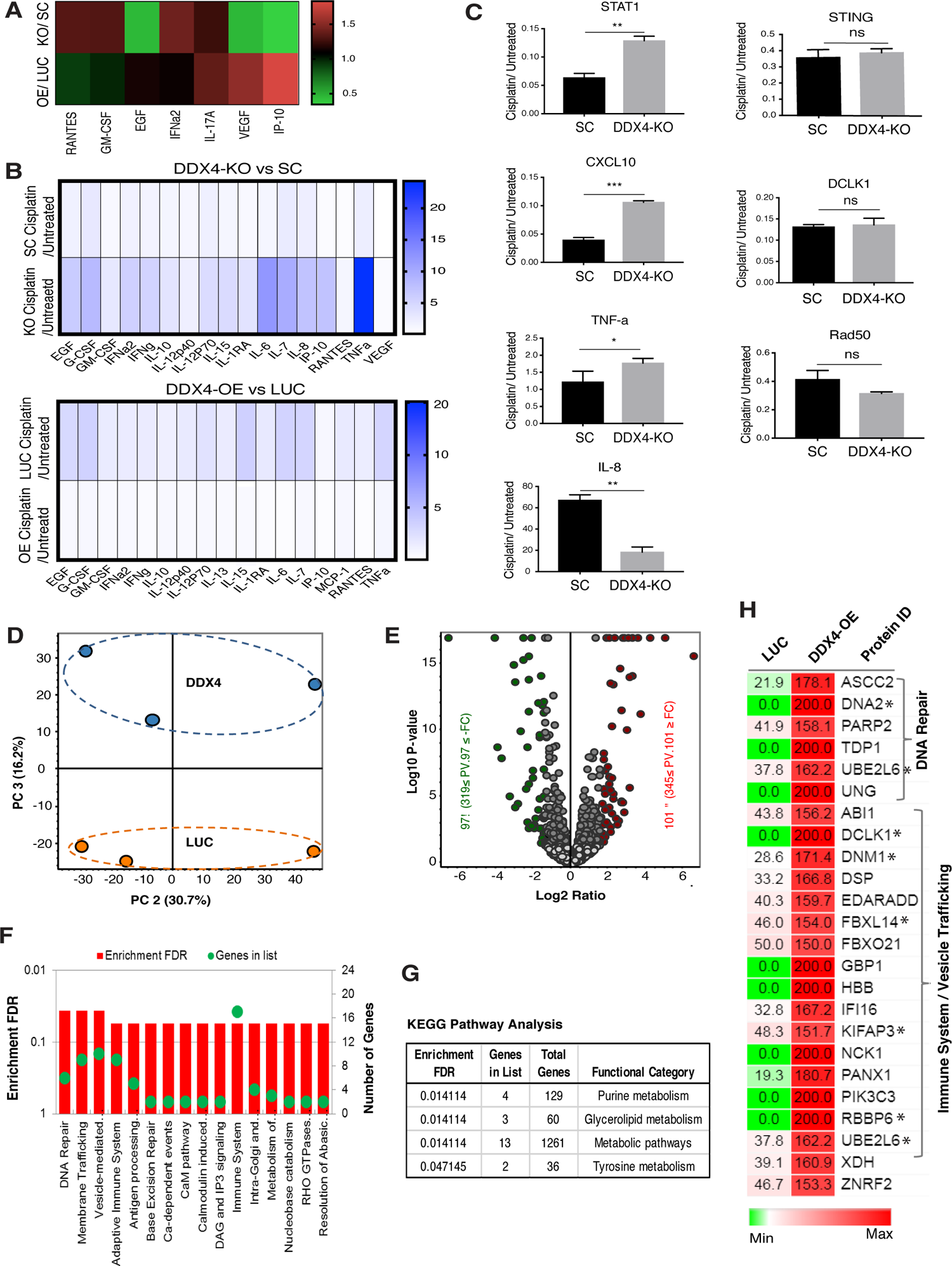
DDX4 globally changes protein expression in H69AR cells. **(A)** Multiplex cytokine profiling shows DDX4-OE has increased cytokine expression compared to the DDX4-KO cell line at a natural state in H69AR cells. Each value was normalized to that of the corresponding control line for each cytokine. **(B)** Multiplex cytokine profiling was performed before and after Cisplatin treatment (10uM Cisplatin treatment for 20 hours) for each cell line of H69AR cells. Each value of the “after” treatment group was normalized to that of the “before” treatment group to obtain the relative expression level of each cytokine after Cisplatin treatment. **(C)** RT-qPCR results of cytokine-related genes. The expression level of each gene was normalized by that of *h36B4*, a housekeeping gene to obtain a relative expression level. The graphs are the average of 2∼4 replicates. **(D-G)** Comparative label-free quantitative proteomic profiling of the DDX4-OE and control (LUC) cell lines. **(D)** Principal component analysis (PCA) of total protein abundance data collected from DDX4 and LUC samples. Data represents the close clustering of protein abundance of each replicates under the same group, however, showed variability between DDX4 and LUC samples. **(E)** Volcano plot of fold change versus q-value of the total of 4740 proteins quantified from DDX4 and LUC cell lines, respectively. Red and green circles represent the significant (q < 0.05) up and down-regulated proteins. Gray circles (q = 0.05) are non-significant and below the threshold of fold expression. **(F-G)** Reactome (F) and KEGG (G) pathway analyses of the significantly increased (red bars) proteins in DDX4-OE vs LUC. **(H)** Heat map analysis of the DNA repair and immune system associated proteins that were significantly increased in abundance in DDX4 OE cell lines. Expression of the proteins with * was validated by immunoblot and/by immunofluorescence.

We next performed label-free quantitative proteomics for DDX4-OE and control (LUC) groups in H69AR cells, which identified over 5000 proteins in each group (Fig. 7D-H; Supplementary Datasheet 1). Gene Ontology (GO) analysis indicates that various pathways were upregulated by DDX4-OE, including DNA/base-excision repair, response to DNA-damage stimulus, protein/macromolecules modifications, polyubiquitination, innate and adaptive immune systems (Fig. 6F). Importantly, all of these factors fell under the category of metabolism by KEGG pathway analysis, suggesting a possibility that DDX4 changes the molecular landscape by controlling the metabolism in the cell (Fig. 6G). These proteomics results were further validated by immunoblot and immunofluorescence for representative proteins using both DDX4-OE and -KO cell lines (Fig. 6H and S7A-B). Similar protein expression trends were also observed in SHP77 (Fig. S7A-B). Further, DDX4-OE cells treated with inhibitors of DNA-damage sensing and immune signaling pathways compromised cellular motility, functionally validating the results of proteomics (Fig. S6C). Taking together, DDX4 appears to be involved in many different biological processes, yet its fundamental function might be in the regulation of metabolism, facilitating cellular fitness and survivability especially under a challenging environment.

**Fig. 7.**
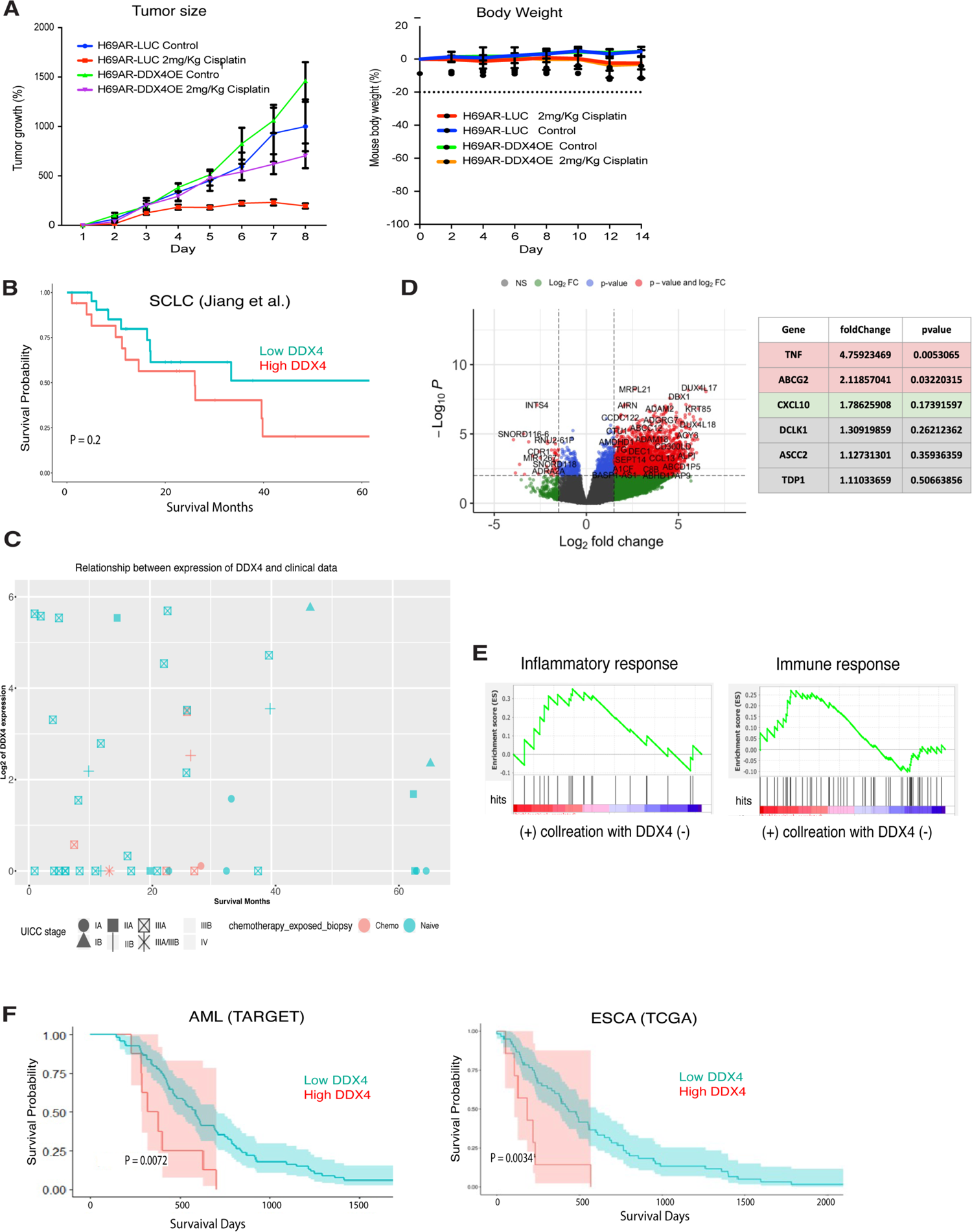
In vivo and clinical implication of DDX4 expression in SCLC and other cancer types. **(A)** LUC (control) or DDX4-OE H69AR cells were injected into five nude mice. (i). DDX4-OE xenografts showed a statistical difference in their size over time after cisplatin treatment compared to the control (LUC) ones. **(B)** Survival SCLC patient dataset. mRNA expression data of the early stage SCLC was obtained from the previously published study of Chinese patients who were diagnosed with SCLC (Jiang et al., 2016). The analysis was limited to pre-treatment samples in case chemotherapy alters DDX4 expression. The SCLC patients with pre-treatment tumors and high DDX4 showed a trend of short survival in the later stage, whereas SCLC patients who survived the longest showed a trend of low DDX4 mRNA expression **(C)** PCA analysis of 40 SCLC samples that are treatment naïve and associated with diagnosis using the dataset available in Jiang et al. (2016). A trend of longer survival for SCLC patients was found with low DDX4 expression (log2 (DDX4) < 2) compared with high DDX4 expression (p-value 0.2). **(D)** Volcano plot of fold change versus p-value of the total of genes quantified from DDX4-High and DDX4-Low patients, respectively. Red, green, blue circles represent the significant (p < 0.01; fold change > 1.5) up and down-regulated genes. Gray circles (p >0.01; fold change < 1.5) are non-significant and below the threshold of fold expression. **(E)** Enrichment analyses of genes for each category of immune response and inflammatory response. Enrichment score indicates the degree of over-representation of each gene set. Red indicates higher rank in the gene set is enriched and blue indicates that lower rank in the gene set is enriched. **(F)** Survival analyses of AML and ESCA patient datasets. mRNA expression datasets were obtained from TARGET or TCGA.

### DDX4 promotes tumorigenesis after chemotherapy

To test *in vivo* the hypothesis that DDX4 facilitates cell survivability under the challenging environment, DDX4-OE or control (LUC) H69AR cells were subcutaneously injected into the nude mice, in the presence or absence of cisplatin, respectively. In the absence of cisplatin, the DDX4-OE tumors showed a similar size and weight compared to control (LUC) tumors (Fig. 7A). In the presence of cisplatin, tumor growth was dramatically reduced in the control (LUC) tumors, but not in the DDX4-OE tumors. The body weight of mice was similar among all groups tested, suggesting that DDX4-OE specifically increased drug resistance and promoted tumorigenesis *in vivo* (Fig. 7A). These results suggest that DDX4 expression facilitates tumor progression even under a challenging environment such as cisplatin treatment, which may also contribute to cancer recurrence after chemotherapy.

### Clinical implication of DDX4 expression in SCLC

To assess the clinical implications of DDX4 expression in SCLC, we analyzed publicly available RNA sequencing results of SCLC patients’ samples. mRNA expression data of early-stage SCLC (Jiang et al., 2016) showed a trend toward worse survival with high DDX4 expression especially at the later stage (Fig. 7B). This suggests that pre-treatment SCLC tumors with high DDX4 expression may be linked to shorter survival in the later stages, while SCLC patients with low DDX4 expression may result in longer survival (Fig. 7B). To test this hypothesis further, we performed survival analysis on 40 samples that were treatment naïve and associated with the survival data. We found a trend of longer survival for SCLC patients with low DDX4 expression compared to the patients with high DDX4 expression (p-value 0.2) (Fig. 7C). This further supports a contention that DDX4 expression may be linked to the survival of SCLC patients especially toward the late stage.

To investigate the potential role of DDX4 in SCLC patients, we performed differential gene expression (DGE) analyses using the same dataset and identified that many genes are upregulated in the DDX4-High patient group (Fig. 7D). Enrichment analyses suggest that genes involved in immune/inflammatory response were found enriched in the DDX4-high patient groups. Since the proteomics results of the cell lines suggest that DDX4 facilitates metabolic protein expressions, these changes in transcript expressions may be the outcome of the DDX4 expression impacting the cellular metabolism in SCLC tumors. Although no proteomics dataset of SCLC patients is currently available, such a dataset will be helpful in the future to identify the direct mRNA targets of DDX4 and how they contribute to patients’ prognosis.

### Implications of DDX4 expressions in somatic cancers

The DGE analysis also reveals that genes related to the germline factors and piRNA biogenesis are enriched in the DDX4-High patient group (Fig. S9A-B). This implies that a cluster of germline factors is indeed expressed in patient tumors as proposed previously (Simpson et al., 2005) and their expressions negatively correlate with patient survivability. What upstream mechanism facilitates DDX4 and other germline factor expressions in somatic cancers is currently unknown and requires further investigation in the near future. To obtain an insight into the clinical implication of DDX4 expression in other somatic cancers, we also performed the survival analyses using publicly available patient datasets of acute myeloma (AML), esophageal carcinoma (ESCA), lung adenocarcinoma (LUAD), bladder carcinoma (BLCA), and uterine carcinoma (UCA) from TCGA or TARGET. Among those, DDX4 expression compromised patients’ survivability in AML and ESCA (Fig. 7F), while it showed no significance or the opposite trend in LUAD, BLCA, and UCA (Fig. S9C). These results suggest that DDX4 expression could contribute to tumorigenesis in both directions. Importantly, the max expression range of DDX4 in SCLC, AML, and ESCA datasets is above 0.06, while that in other cancers is much below 0.6 or 0 (Fig. S9C). This implies that DDX4 expression facilitates tumorigenesis only in certain cell types that are equipped to tolerate its higher expression, while other cell types may not tolerate DDX4 expression and die immediately. This makes sense because Vasa/DDX4 expression in somatic cells is tightly regulated under normal circumstances and its ectopic expression is known to cause developmental failure in embryonic cells (Fernandez-Nicholas et al., *in press*). This hypothesis can be further tested in the future using cancer cell lines that do not express DDX4. Identifying how the forced expression of DDX4 changes the molecular expression profiles in each cancer cell type will help us understand what molecular backgrounds are tolerant or intolerant to DDX4 expression and contribute to tumorigenesis or cell death in each cancer type, respectively.

### Conclusions: DDX4 functions in somatic cancers

DDX4/Vasa knockdown often results in infertility in various organisms. Therefore, it had been conventionally used exclusively as a germline marker and implicated in mRNA translation of germline-related factors (Hay et al., 1988; Lasko and Ashburner, 1988; Linder, 2006; Raz, 2000; Sengoku et al., 2006; Gustafson and Wessel, 2010; Lasko, 2013; Liu et al., 2009). In this study, however, we report essential functions of DDX4 outside of the germline *in vitro, in vivo,* and in clinical datasets. Since DDX4 expression in cancer cells is relatively low compared to other oncogenic drivers, dramatic phenotypes caused by DDX4-OE or DDX4-depletion in SCLC cells were initially surprising. This suggests that DDX4 may not act as an oncogene, but instead promote chemoresistance in certain cancer cells, contributing to cancer recurrence after chemotherapy.

Indeed, the past report in epithelial ovarian cancer patient tissues suggests that DDX4 is consistently co-expressed with a stem cell marker CD133 and only in a small fraction of cells within a tumor (Kim et al., 2014). Future studies of DDX4 immunohistochemistry in various patient tissues will be important to test this possibility.

The detailed molecular mechanisms of DDX4 expression outside of the germline also remain unclear. Based on the findings in this study, DDX4 appears to regulate metabolic protein expressions, facilitating changes in the molecular landscape of the cell, especially in a challenging environment. This model explains the critical importance of DDX4 expression, even at low levels. Further, DDX4 is heavily regulated by post-transcriptional and post-translational modifications, and a low transcript or protein level may not directly imply low DDX4 function. This was indeed reported to be the case during brain tumor induction in *Drosophila* (Janic et al., 2010). Further mechanistic studies with animal models are needed to understand the mechanisms of DDX4 function in cancers.

Lastly, since DDX4 is generally not expressed in adult somatic cells and (Poon et al., 2016), thus it may represent a potential therapeutic target in DDX4 expressing cancers that are likely highly resistant. Although germ cells may be also affected, the importance of these side effects will vary depending on the patient’s life stage. The DDX4-C3 mutant which we identified as critical for DDX4’s function in SCLC cells is highly specific to DDX4 but not conserved in other DEAD-box family proteins. Therefore, this region has the potential to serve as a promising target for small molecule inhibitors in the future. Conversely, the DDX4 expression may be engineered to eliminate certain tumorigenic cells that have no tolerance to DDX4 expression. Further investigations in these directions will be useful to evaluate DDX4’s potential as a new therapeutic target.

## Author Contributions

C.N. and S.K. were responsible for the concept, experimental design and undertaking, data analysis, and manuscript construction and editing; X.L. was responsible for experimental design and undertaking, data analysis, and manuscript editing related to the in vivo mouse experiments; A.H, E.K., Y.S., S.M., and N. T. were responsible for experimental design, undertaking, data analysis, and manuscript construction related to the clinical sample experiments. N.A. was responsible for experimental undertaking and analyses of the proteomics dataset. J.M. and T.T. were responsible for the experimental design and undertaking of the CTG assay. T.T., T.X., and D.X were responsible for cell line construction, cell culture, and cell motility assays; N.T., C.T., and D.B. were responsible for experimental design and manuscript editing; M.Y. was responsible for concepts, experimental design, and undertaking, data analysis, manuscript construction, and editing.

## Acknowledgments

We thank the members of the Cancer Center at Brown University for active discussions.

## Data Availability

The datasets used and/or analyzed during the current study are available from the corresponding author on reasonable request.

## Consent for publication

No individual person’s data is contained within the manuscript.

## Ethics approval and consent to participate

No primary patient data is used in this study.

## SUPPLEMENTARY FIGURES & LEGENDS & TABLES

**Fig. S1.**
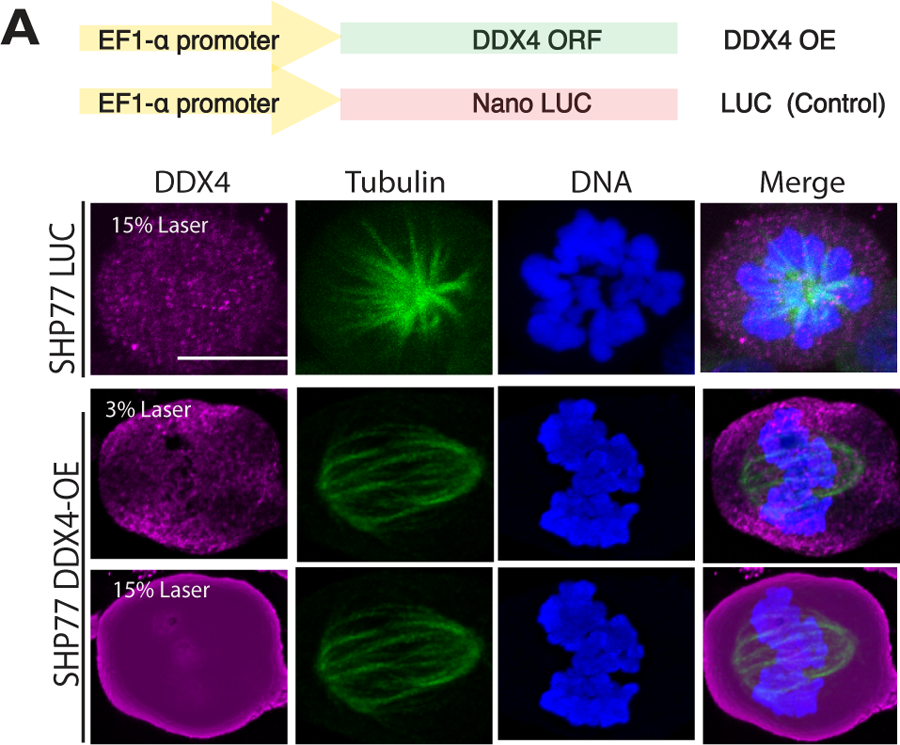
DDX4 overexpression in SHP77. DDX4-ORF was driven by the EF1-α promoter (upper diagram), which showed overexpression of DDX4 protein (magenta) by immunofluorescence in SHP77 cells. Tubulin, green; DNA, blue. Scale bar = 5 μm.

**Fig. S2.**
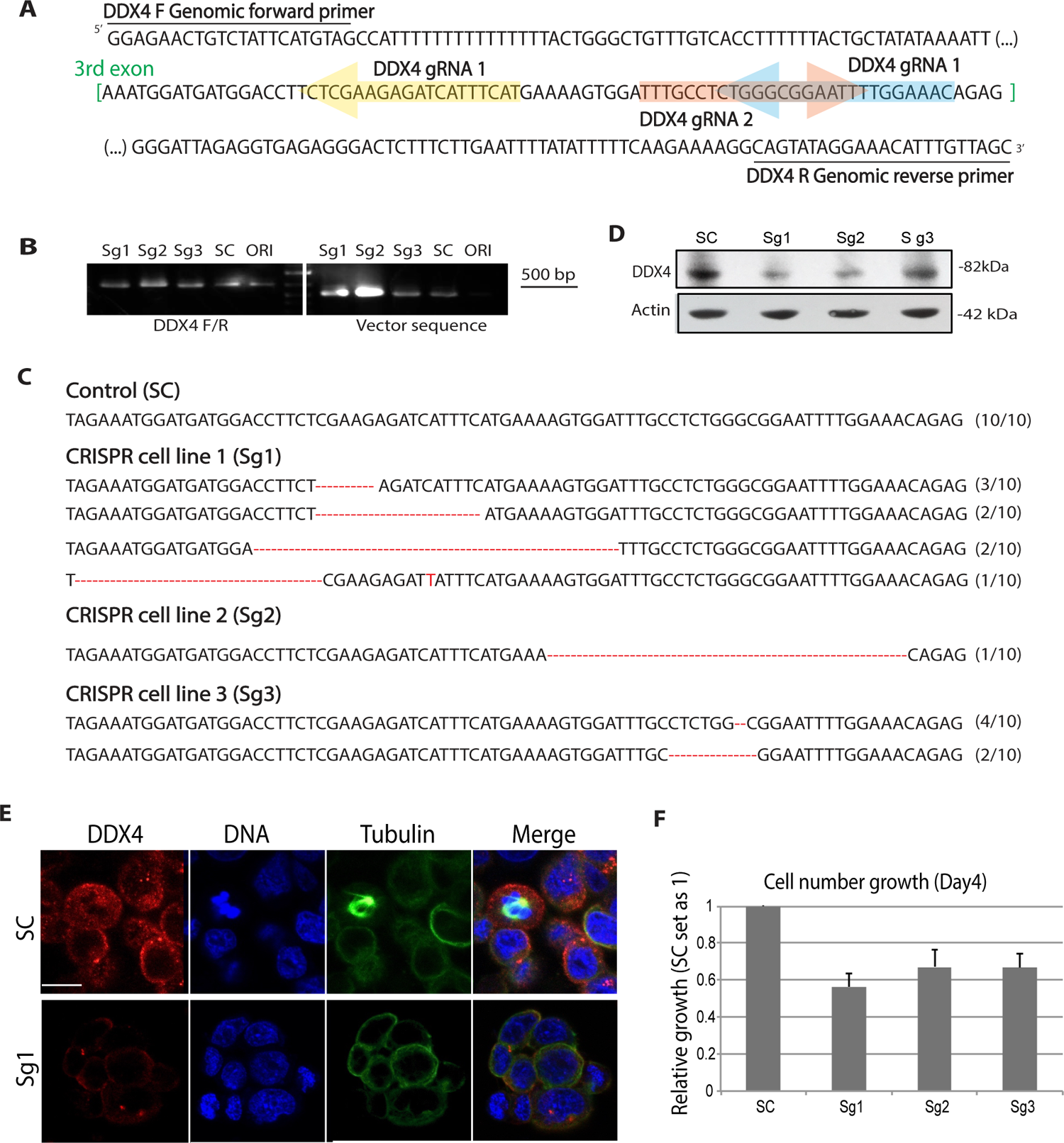
DDX4 knockout cell construction in H69AR by CRISPR-Cas9 gene-editing technology. (A) Three different gRNAs were designed within the 3rd exon of the DDX4 gene to construct knockout cell lines (Sg1-Sg3). (B) Left, Genomic PCR results of the DDX4 genomic locus shown in A for three CRISPR knockout cell lines (Sg1-Sg3), control cells introduced with scrambled gRNA sequence (SC), and parental intact cells (ORI). Right, Genomic PCR results detect the lentiviral vector sequence used for virus infection, showing the effective introduction of CRISPR constructs in each cell line. (C) DNA sequencing results of the 3^rd^ exon and its flanking region of the DDX4 gene depicted in A. The Sg1 cell line showed deletions in many of the clones sequenced compared to other KO lines, and the control (SC) line showed no mutations. () in the right corner suggests the number of the genomic PCR clones showing the corresponding sequence. (D-E) Immunoblot (D) and Immunofluorescence (E) results showed reduced DDX4 protein expression in Sg1 and Sg2 knockout cell lines compared to control (SC). Actin was used as a loading standard for Immunoblot. (F) The cell number growth comparison among DDX4-KO cell lines. The same number (1×10^5^) of cells was harvested for each cell line and the total number of cells was counted on Day 4. Each value was normalized to that of the control line (SC). The results are the average of three independent experiments.

**Fig. S3.**
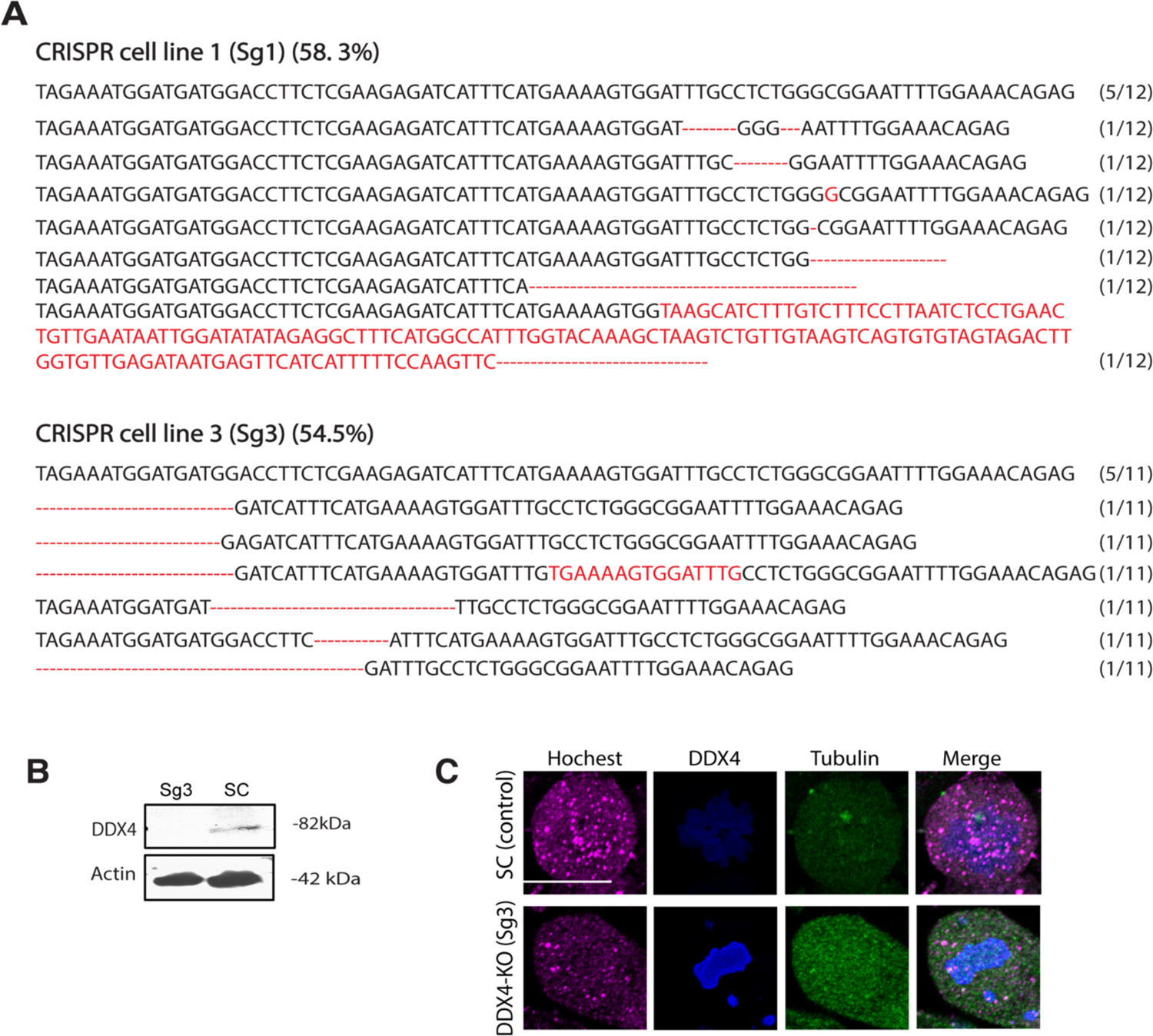
DDX4 knockout cell construction in SHP77. (A) DDX4-KO was constructed by CRISPR-Cas9 gene-editing technology. The same lentivirus (Sg1 & Sg2) used in Fig. S1 for H69AR cells were used in SHP77 cells. DNA sequencing results of the 3^rd^ exon and its flanking region of the DDX4 gene depicted in A. The Sg3 cell line showed deletions in many of the clones sequenced compared to other KO lines, and the control (SC) line showed no mutations. () in the right corner suggests the number of the genomic PCR clones showing the corresponding sequence. (B-C) Immunoblot (B) and immunofluorescence (C) results showed reduced DDX4 protein expression (magenta) in Sg3 knockout cell lines compared to control (SC). Tubulin, green; DNA, blue. Scale bar = 5 μm.

**Fig. S4.**
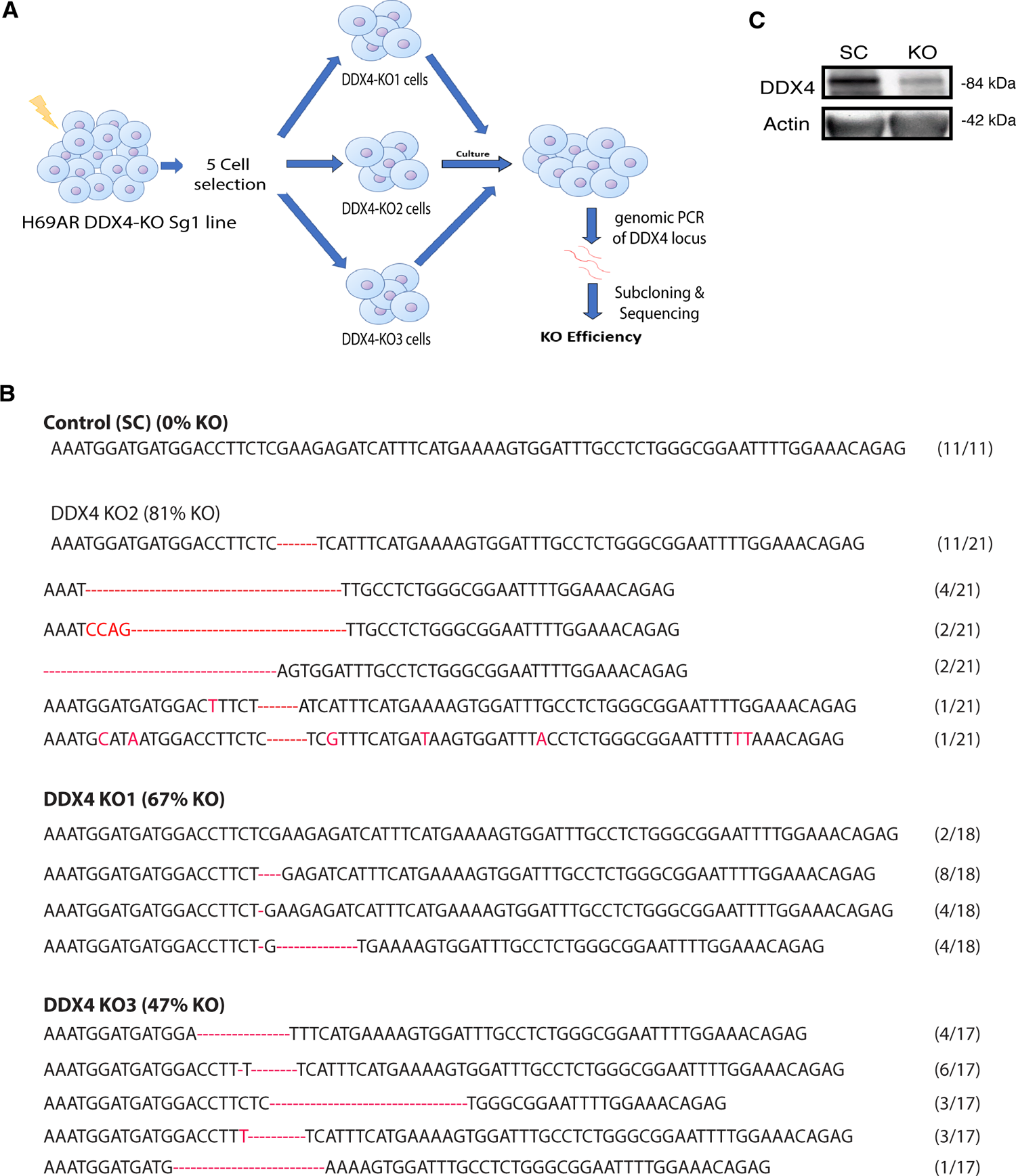
Five cell selection of the DDX4-KO lines in H69AR cells. (A) A schema showing the process for the five-cell selection of DDX4-KO cells made in Fig. S1. The five groups were selected for DDX4-KO Sg1 and SC lines, respectively. Since only three out of five groups of DDX4-KO survived, the genomic PCR analysis was performed for these three groups (KO1, KO2, KO3). (B) KO2 exhibited the greatest DDX4-KO efficiency by inducing a stop codon either within or immediately after the 3^rd^ exon of DDX4, whereas the same five-cell selection of SC lines resulted in no mutation. () in the right corner suggests the number of the genomic PCR clones showing the corresponding sequence. (C) Immunoblot results of SC (control) and KO (DDX4-KO) cells. Actin was used as a loading standard.

**Fig. S5.**
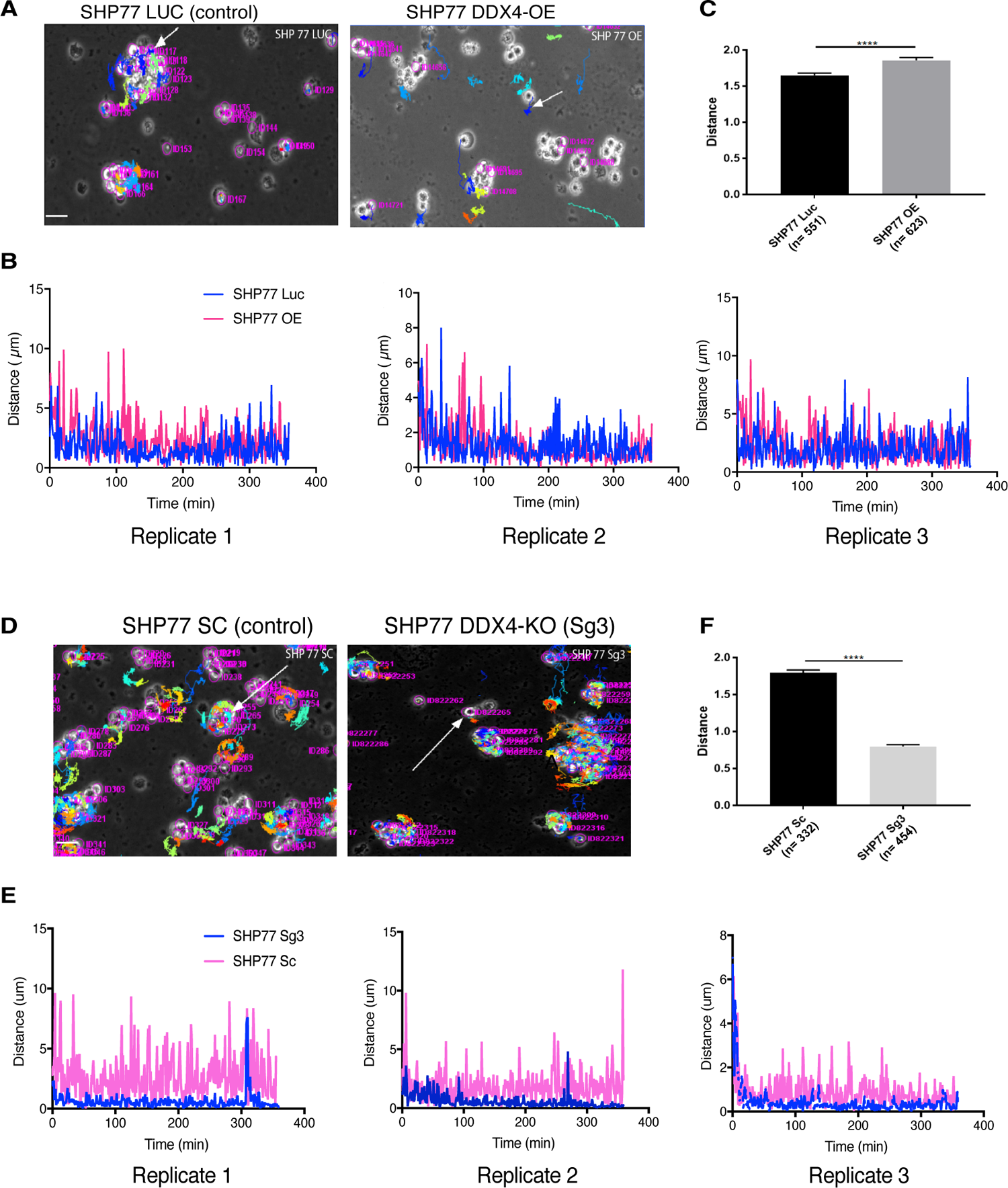
DDX4 increased cell motility in SHP77 cells. **(A-C)** Each of the four cell lines (DDX4-OE, LUC, DDX4-KO, SC) was timelapse-imaged for 6∼12 hours, with a new image being taken every minute. The representative images of the paths (colored lines) taken by the tracked cells are shown. The travel distances of the cells pointed to by the arrows were compared among cell lines and shown for each cell over the time point (Aʹ). **(D-F)** SHP77 SC (control for KO) and OE showed greater motility than SHP77 KO (Sg3) and LUC (control for OE), respectively. **(C & F)** The travel distances of four tracked SHP77 cells per cell line were summated and averaged. DDX4-KO (Sg3) showed lower average motility and DDX4-OE was a higher one when compared to controls. () indicates the total number of cells analyzed. Scale bar = 10 μm.

**Fig. S6.**
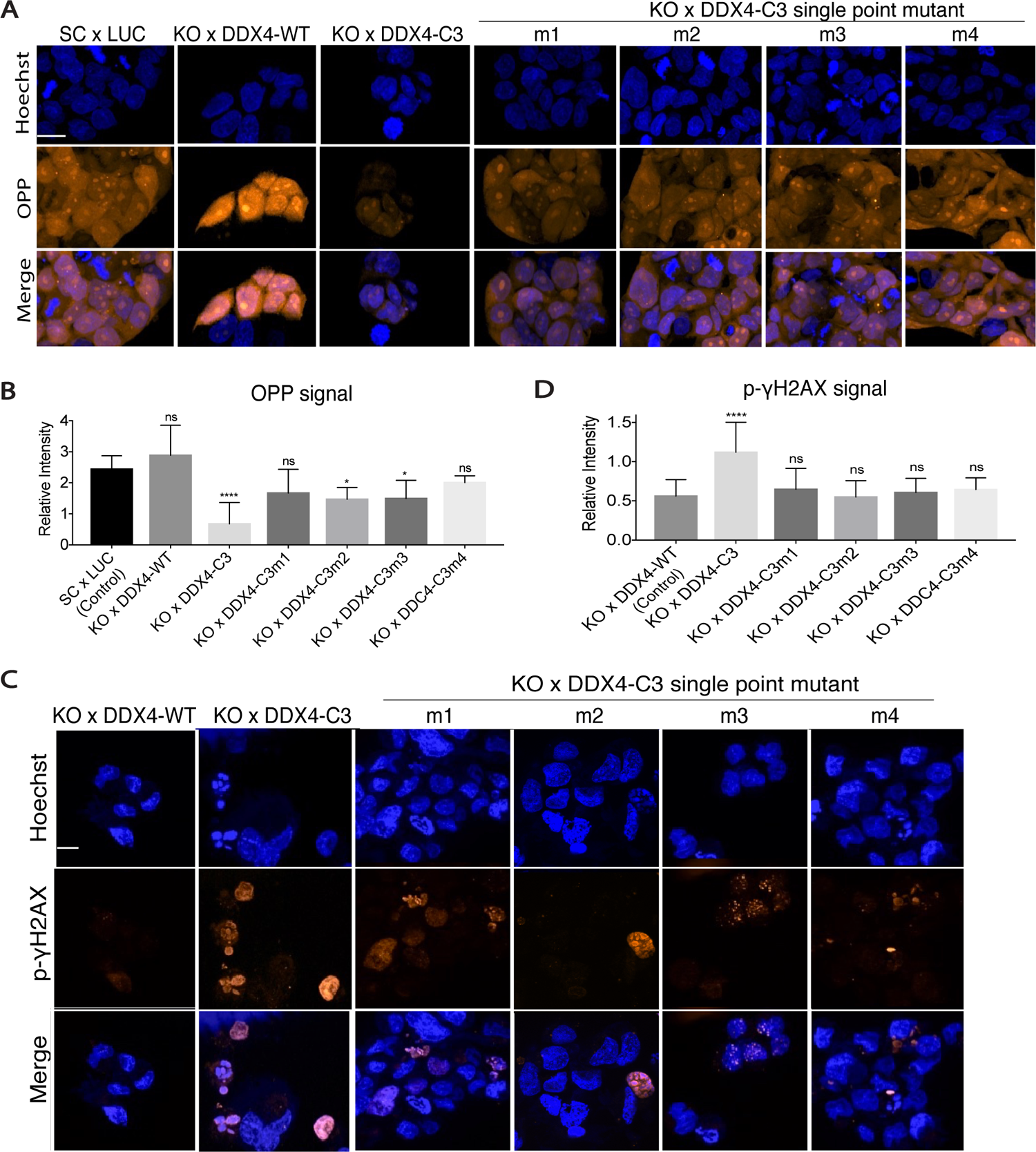
A single point mutation in the C3 region of DDX4 rescued the DDX4-KO phenotypes in H69AR cells. Translational activity (A-B) or DNA-damage level (C-D) was measured by the OPP signal or by the p-γH2AX signal level for each sample group, respectively. The OPP or p-γH2AX signal intensity was normalized to the Hochest signal level in the same ROI for each image. 8 ROIs were measured per sample group and the average value of each group is shown in the graph. Scale bars = 10 μm.

**Fig. S7.**
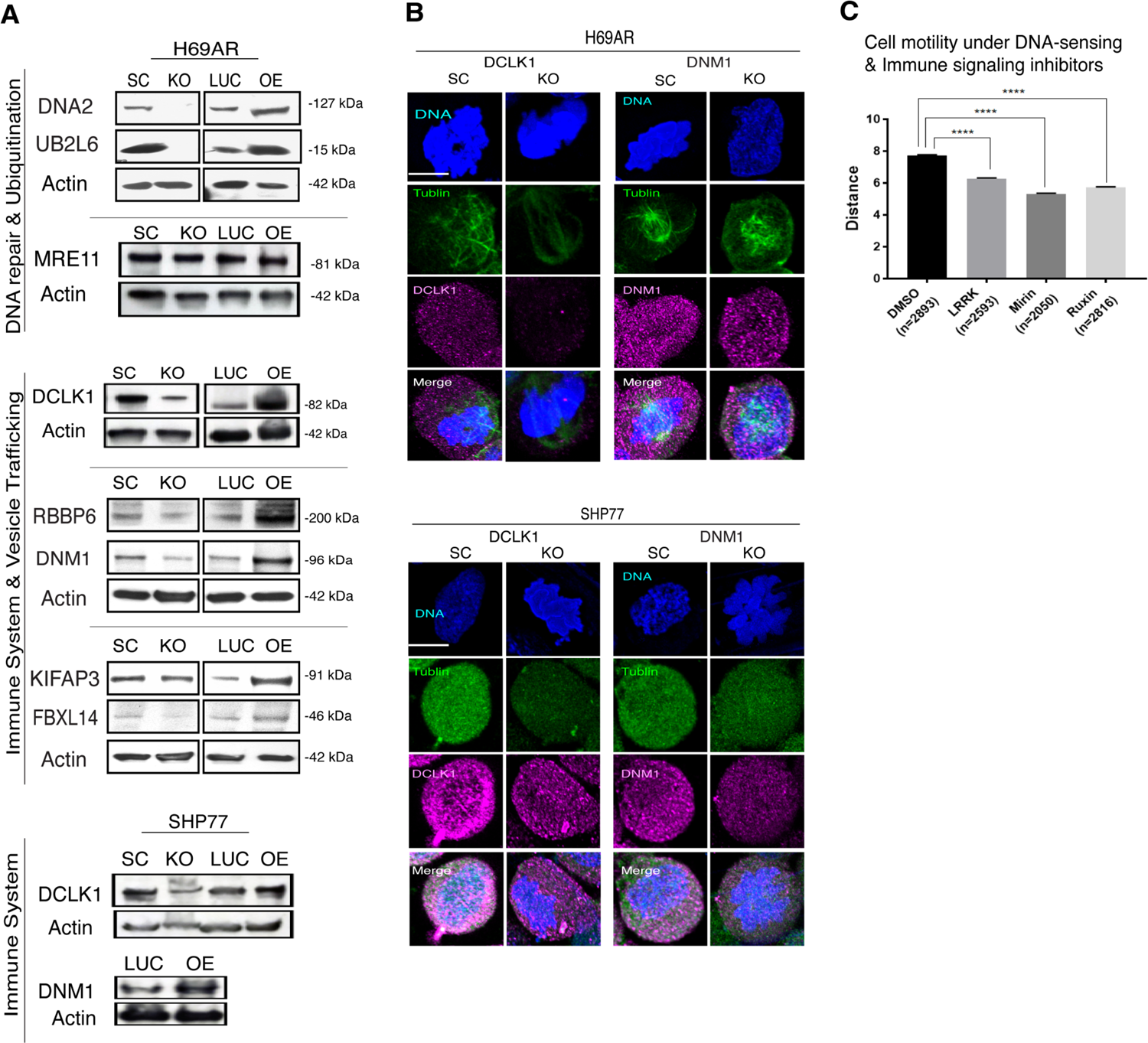
DDX4 expression upregulated the metabolic pathways in H69AR and SHP77 cells. **(A-B)** Validation of the proteomic results by immunoblot (A) or by immunofluorescence (B). DCLK1, DNA2, UB2L6, DCLK1, RBBP6, DNM1, KIFAP3, FBXL14 levels were upregulated or downregulated by DDX4-OE or –KO, respectively, of H69AR or SHP77 cells. On the other hand, little difference was seen in the expression level of MRE11 (negative control) between DDX4-OE and LUC lines, which is consistent with the proteomics results. Actin was used as a standard for all immunoblots. Scale bars = 5 μm. DNA, blue; Tubulin, green. **(C)** Inhibitors for immune response (LRRK and Ruxin) and DNA-damage sensing (Mirin) all reduced cell motility in DDX4-OE cells.

**Fig. S8.**
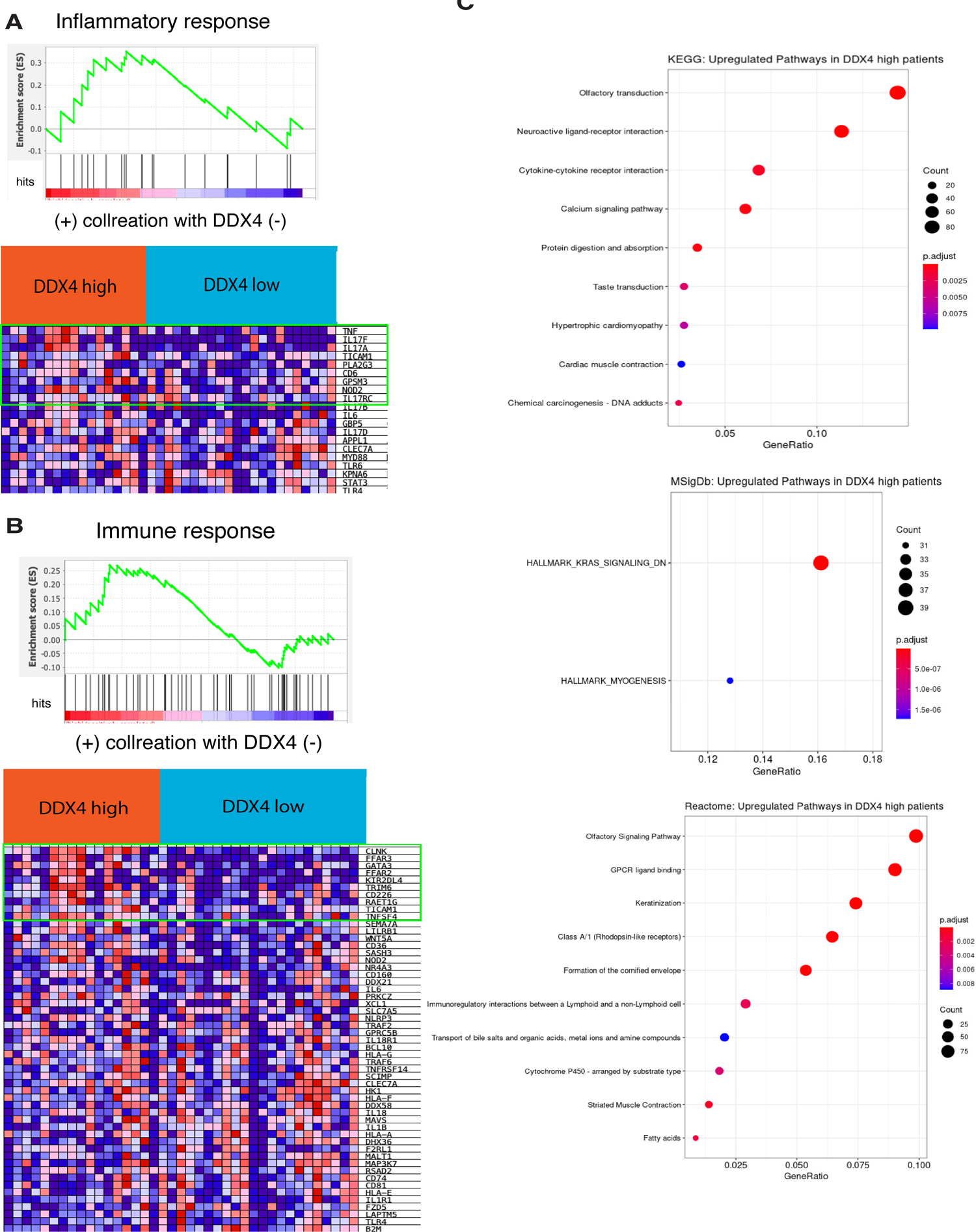
Functional analysis and heat-maps for enriched genes of each category in the DDXD4-High SCLC patients, linked to Fig. 6E. (A) Inflammatory response-related genes. (B) Immune-response-related genes. (C) Functional analysis suggests DNA abuducts, metabolism and K-ras signaling are enriched in DDX4-high SCLC patients.

**Fig. S9.**
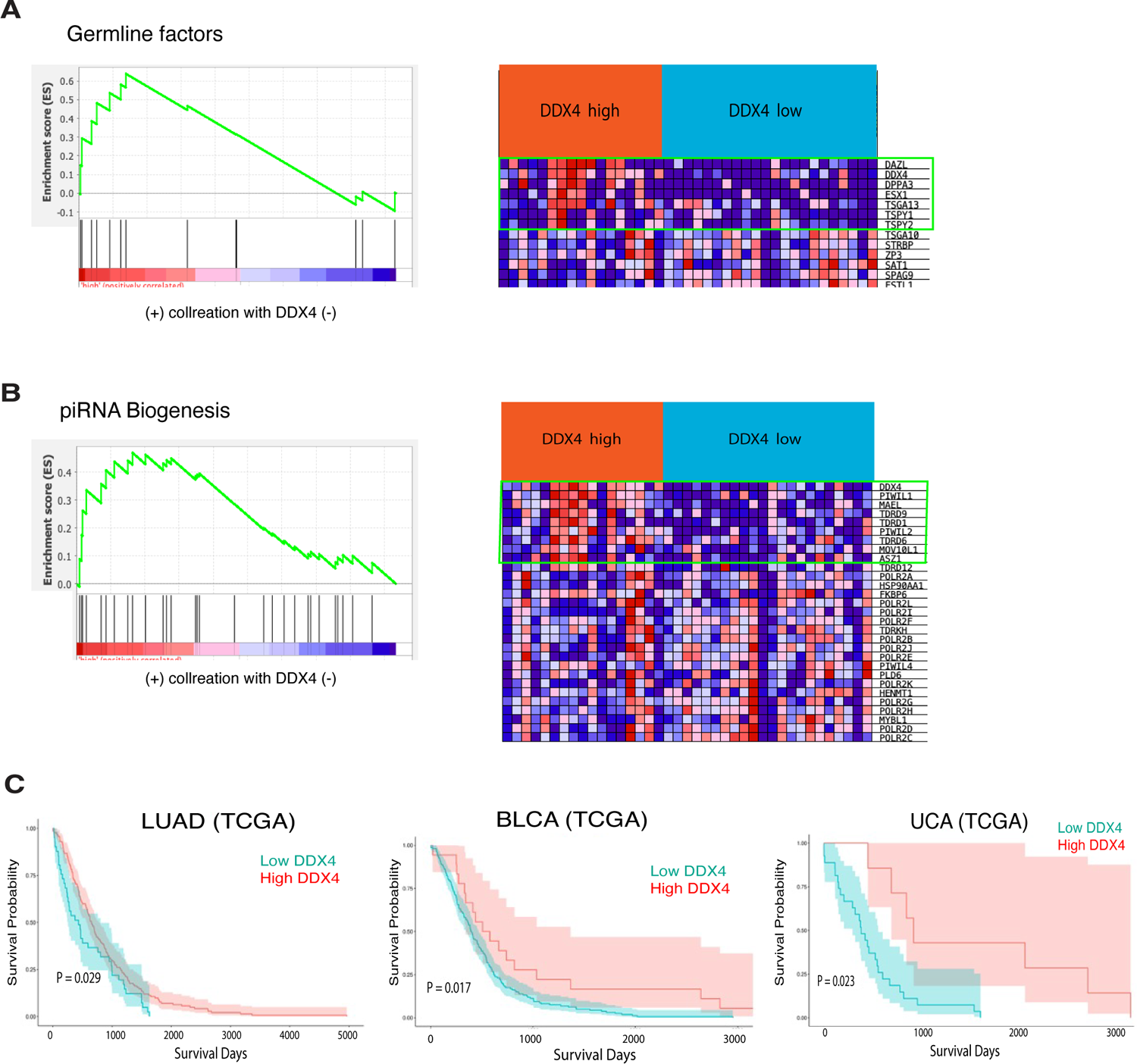
Enrichment analyses and corresponding heatmaps, and survival analyses, linked to Fig. 6E. (A-B) Enrichment analyses for germline factors (A) and piRNA biogenesis molecules (B) in DDX4-High SCLC patients. (C) Survival analyses. mRNA expression datasets of LUAD, BLCA, and UCA patients’ samples were all obtained from TCGA.

## SUPPLEMENTARY MOVIE LEGENDS

**M1-M4.** Each of LUC(M1), DDX4-OE (M2), SC(M3), and DDX4-KO (M4) H69A cell-line was time-lapse imaged at 30 minutes intervals for 24∼48 hours.

## Methods

### Outlines

#### KEY RESOURCES TABLE

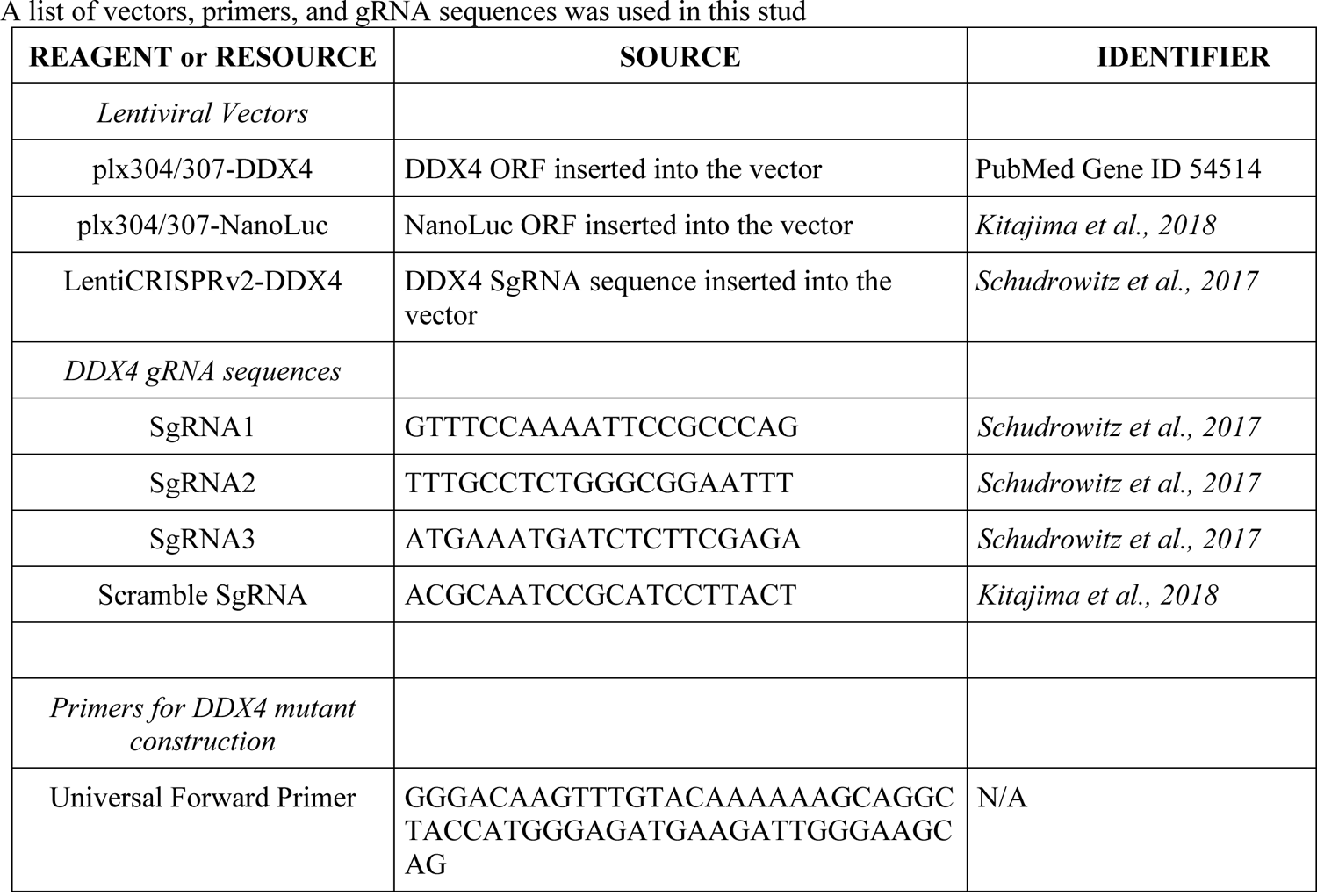
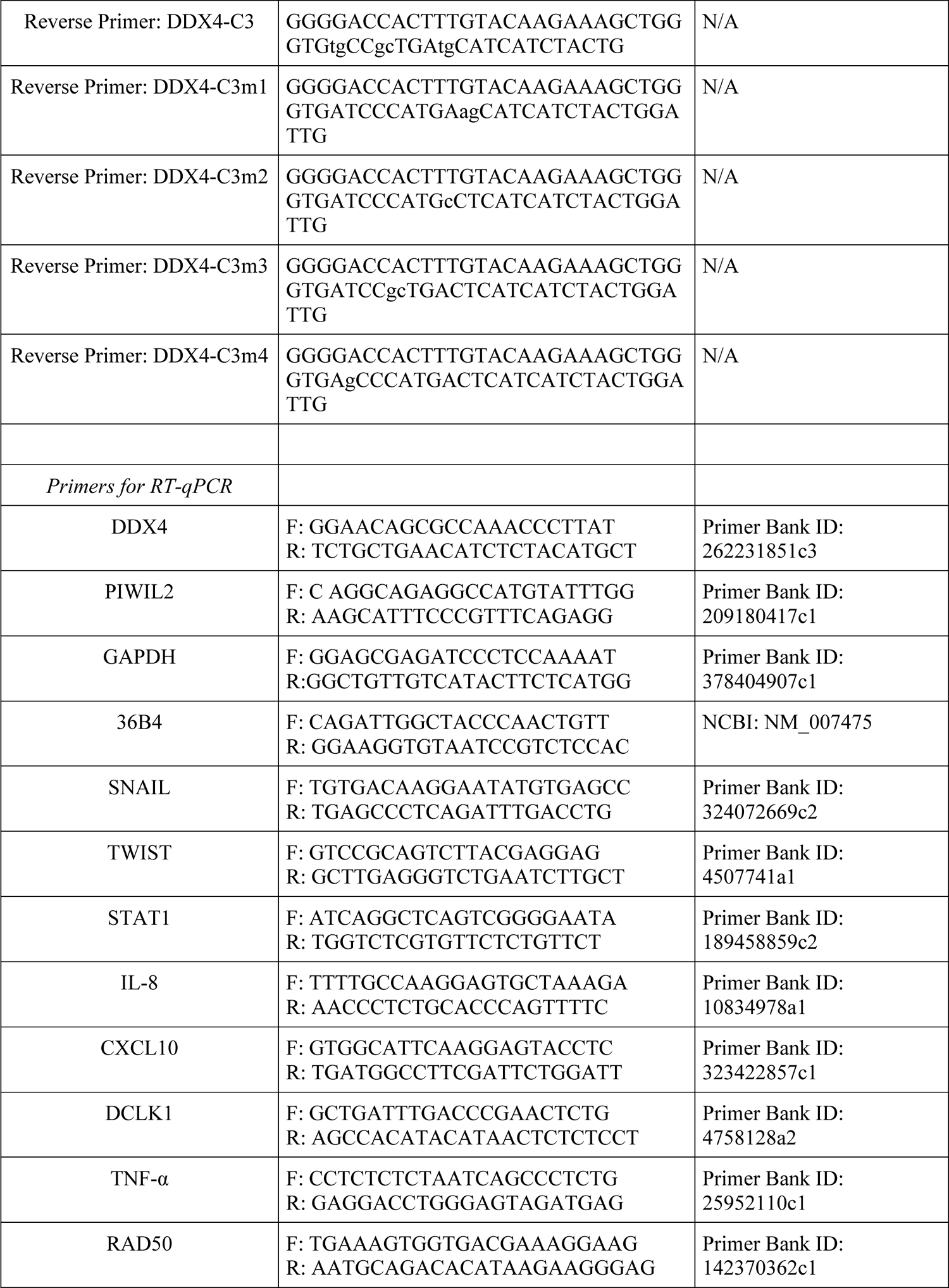
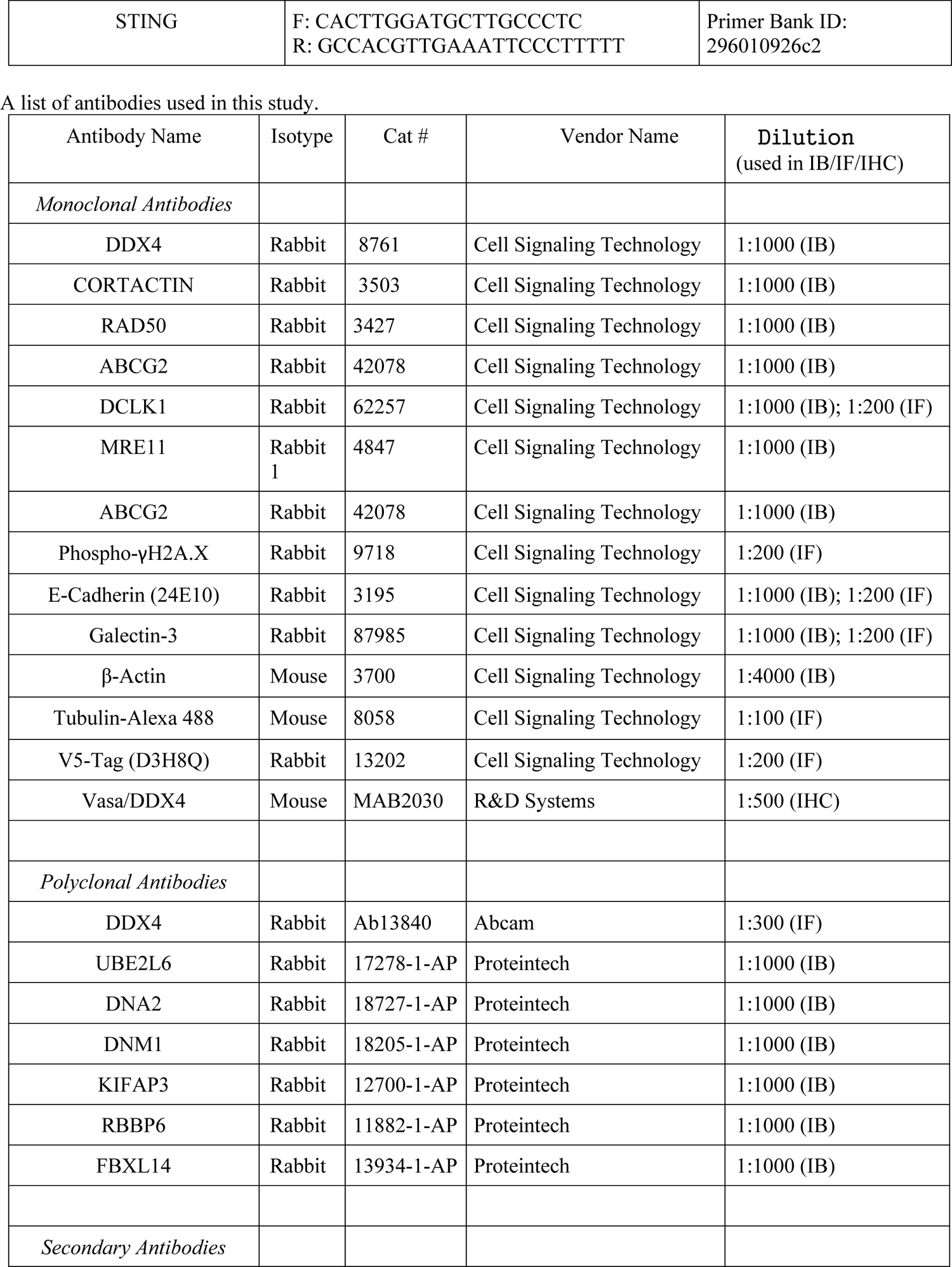
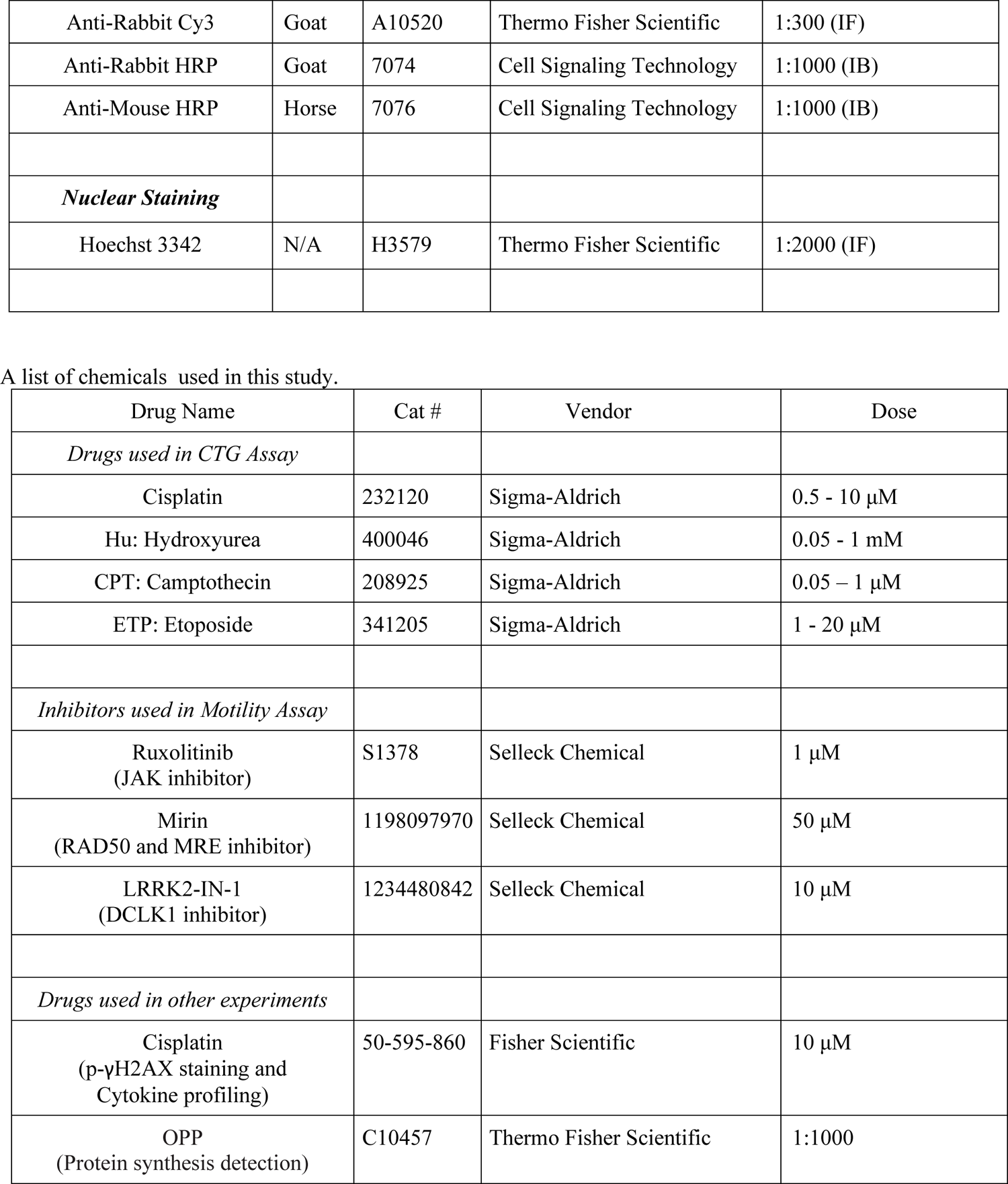

##### CONTACT FOR REAGENT AND RESOURCE SHARING

For further information, requests should be directed to and will be fulfilled by the Lead Contact, Mamiko Yajima.

##### EXPERIMENTAL MODEL AND SUBJECT DETAILS

All human cancer cell lines were obtained from ATCC. Mouse studies were performed in accordance with guidelines of the Institutional Animal Care and Use Committee at Kanazawa University. The procedures of cell injection and analysis of tumor growth were described previously (Kitajima et al. Cancer Cell 2018) using KSN/Slc mice (Japan SLC, Inc.).

##### Methods Details

###### Cell lines, cell culture and cell number counting

SCLC cell lines H69AR (ATCC# CRL-11351) and SHP77 (ATCC# CRL-2195), H69 (ATCC# HTP-119), H187 (ATCC# CRL-5804) and H82 (ATCC# HTB-175), as well as Non-SCLC (NSCLC) cell lines H1792 (ATCC# CRL-5895) and H23 (ATCC# CRL-5800) were cultured in RPMI-1640 (#A1049-01, GIBCO) supplemented with 10% (v/v) heat-inactivated fetal bovine serum (ThermoFisher Scientific), and antibiotics, in a humidified atmosphere of 95% air and 5% CO_2_ at 37°C. For all experiments, cells in the log phase of growth were used. For counting cell number, each cell line was suspended at 1x 10^5^ /mL and counted on a hemocytometer every 3 days of culture for five independent times.

###### Fluorescence activated cell sorting (FACS) for five cell selection

A five-colony selection of DDX4-knockout (KO) cell line was performed by FACS sorting (FACS Aria II, BD Biosciences). Five cell each of isolated cells were cultured in each well of the 96 or 24-well plate for ∼2 months and subjected to genomic PCR and sequencing to identify the efficiency of KO.

###### DDX4 overexpression in H69AR and SHP77 cells

Lentiviral vectors for DDX4, DDX4-C3 and NanoLuc (control) overexpression were constructed by inserting DDX4 open reading frame (ORF), DDX4-C3 or NanoLuc sequence into the plx307 or plx304 vector that contains EF1-α promoter at its 5’ end using the Gateway cloning protocol (Thermo Fisher Scientific). DDX4-C3 and C3m1∼m4 mutants were constructed by PCR-amplifying the DDX4-ORF fragment using a primer set containing the mutations to convert all or one of the last three amino acids of <u>E</u>-S-<u>W</u>-<u>D</u> to Alanine (see the resource table for details). Lentiviral infection was performed as described in Shalem et al. (2014) and Kitajima et al. (2019) followed by a puromycin and/or blasticidin selection at the concentration of 2∼10 μg/mL.

###### DDX4 CRISPR-mediated knockout in H69AR and SHP77 cells

Lentiviral vectors for CRISPR-mediated DDX4 knockout were previously constructed and used in myeloma IM9 cells, which resulted in a successful knockout of DDX4 (Schudrowitz et al., 2017). Briefly, following the protocol described in Shalem et al. (2014) and Sanjana et al. (2014), three guide RNAs (gRNAs) were designed within the 3^rd^ exon of DDX4 gene locus (see also the resources table). A scrambled gRNA sequence formed by random combination of A,G,T,C, and which does not share identity with either the mouse or human genome, was used as a control (Kitajima et al., 2019). The efficiency of genome editing in each cell line was analyzed by genomic PCR and sequencing (Fig. S2B), confirming mutations in the 3rd exon of the DDX4 genomic locus in 80% of H69AR Sg1 cell lines, 10% of Sg2, and 60% of Sg3 cell lines (n=10 each) (Fig. S2C). In SHP77, those mutations were less efficient and 42% in Sg1 and 46% in Sg3 (n=10 each) (Fig. S3A). In both cell lines, no mutation was ever found in control SC cell lines (n=11 each). Immunoblot and immunofluorescent results confirmed that these genomic mutations reduced DDX4 protein expression in the Sg1 and Sg3 lines compared to the control cell line SC (Figs. S2D-E and S3B-C).

###### Genomic extraction and genomic PCR

1 μl of each cell pellet was subjected to genomic DNA extraction by treating with 20 μL of QuickExtract DNA Extraction Solution (# QE09050, Epicenter, USA) at 65°C for 6 minutes followed by heat inactivation at 98°C for 2 minutes. 1 μl each of these treated samples was used for genomic PCR to amplify a flanking region of the 3^rd^ exon of DDX4, where the gRNA sequences were designed, using high fidelity HIFI PCR premix (Clonetech, USA) with primers summarized in the resource table. The resultant PCR products were either separated by agarose gel electrophoresis and visualized or subjected to sub-cloning into the Zero Blunt TOPO PCR Cloning Vector (Invitrogen, USA) for sequencing.

###### Reverse Transcription and Quantitative real-time PCR (RT-qPCR)

RNA extraction was performed using RNeasy Mini Kit (QIAGEN, Hilden, Germany), and 1 μg each of the resultant total RNAs were subjected to Reverse Transcription (RT) using the Maxima First Strand cDNA Synthesis Kit (Thermo Fischer Scientific, Waltham, MA) by following the manufacturer’s protocol. 1 μl each of the cDNAs was then used either for conventional PCR reactions or for qPCR reactions. For conventional PCR reactions were performed with 35 cycles using HIFI PCR premix (Clonetech, USA). qPCR was performed using 1 μL of cDNA with Luna Universal qPCR Master Mix (# M3003S, New England Biolabs, USA) following the manufacturer’s protocol. The primers are summarized in the resource table.

###### Immunofluorescence, OPP staining, and microscopy

Cells were fixed with 4% PFA at 4°C overnight or 30 minutes at room temperature, washed with PBS three times, and stained with a primary antibody of interest (see the resource table for details) at 1:100∼1:300 dilution at room temperature for 3∼5 hours. These samples were washed 6 times with PBS and stained with a secondary antibody against rabbit IgG (1:500, 1mg/mL, Invitrogen) in PBS. They were then washed four times and a Hoechst nuclear stain (10 mg/mL, Promega) in PBS was applied at a 1:1500 dilution. For OPP stainig, cells were incubated with O-propargyl-puromycin (OPP, ThermoFisher, # C10457) at 1:1000 dilution prior to fixation and subjected to Click-iT reaction by following the manufacturer’s protocol. The resultant cells were imaged by confocal laser microscopy (Olympus FV3000 and Olympus Spinning disk). Imaging was conducted using the same laser condition throughout each cycle of the experiments with both the control and experimental groups for quantitative accuracy. The signal intensity of each image was measured by *Image J* for quantitative analysis.

###### Immunoblot analysis

Immunoblots were performed by collecting approximately 10 μL of packed cells from each group in 50 μL of loading buffer. Each sample (2 μL) was run on a 10% Tris–glycine polyacrylamide gel and transferred to nitrocellulose membranes for immunoblotting with each primary antibody of interest at 1:1000 (see the resource table for details), and then with peroxidase-conjugated anti-mouse or -rabbit secondary antibodies at 1: 2000 (Cell Signaling Technologies), respectively. The bound antibodies were detected by incubation in a chemiluminescence solution (1.25 mM luminol, 68 μM coumeric acid, 0.0093% hydrogen peroxide and 0.1 M Tris pH 8.6) for 1∼10 minutes, exposed to film and developed. Each experiment was performed at least three independent times. The signal intensity of each band was measured by *Image J* to construct graphs.

###### Cell motility assay

The cells (1×10^5^ cells/ml) from each group were cultured in each well of the 24-well plate and imaged every 30 minutes for H69AR cells or every minute for SHP77 cells in live up to desired time point. Images were taken with 10X objective on EVOS cell imaging microscope with a CO_2_ incubator (Thermo Fisher Scientific). For inhibitor treatment, each inhibitor at the desired dose (see the resources table for details) was added at the beginning of imaging for 24 hours. Analyses were all done using *FIJI’s trackmate*. Cell motility was analyzed by measuring the position of cell centroid at every time point and plotted to show the trace of centroid movement. The distance that the cell centroid traversed for each time point was calculated to determine the level of the movement.

###### Drug treatment and Cell survivability (CTG) Assay

For drug treatment, details are listed in the resource table or described in the text for each experiment. For cell survivability assay, each sample group (1×10^5^ cells/ml) were harvested in each well of the 96-well plate, treated with a drug of interest or with DMSO (for control) with various concentrations for three days, and subjected to Cell Titer-Glo Luminescent Cell Viability (CTG) Assay (Promega, USA) by following the manufacturer’s protocol. The data was normalized by the value of the control (DMSO-treated) group.

###### Quantitative proteomics and Data analysis

For each cell lines [LUC (Control) and DDX4-OE (Experimental)] 4×10^6^ H69AR cells were lysed with a lysis buffer (8 M urea, 1 mM sodium orthovanadate, 20 mM HEPES, 2.5 mM sodium pyrophosphate, 1 mM β-glycerophosphate, pH 8.0, 20 min, 4°C) followed by sonication at 40% amplification by using a microtip sonicator (QSonica, LLC, Model no. Q55) and cleared by centrifugation (14 000 × g, 15 min, 15°C). Protein concentration was measured (Pierce BCA Protein Assay, Thermo Fisher Scientific, IL, USA) and a total of 100 µg of protein per sample was subjected for trypsin digestion. Typtic peptides were desalted using C18 Sep-Pak plus cartridges (Waters, Milford, MA) and were lyophilized for 48 hours to dryness. The dried eluted peptides were reconstituted in buffer A (0.1 M acetic acid) at a concentration of 1 µg/µl and 5 µl was injected for each analysis.

The LC-MS/MS was performed on a fully automated proteomic technology platform (Ahsan et al., 2017). that includes an Agilent 1200 Series Quaternary HPLC system (Agilent Technologies, Santa Clara, CA) connected to a Q Exactive Plus mass spectrometer (Thermo Fisher Scientific, Waltham, MA). The LC-MS/MS set up was used as described earlier (Ahsan et al., 2017, J Proteomics 2017, 165: 69-74). Briefly, the peptides were separated through a linear reversed-phase 90 min gradient from 0% to 40% buffer B (0.1 M acetic acid in acetonitrile) at a flow rate of 3 µl /min through a 3 µm 20 cm C18 column (OD/ID 360/75, Tip 8 µm, New objectives, Woburn, MA) for a total of 90 min run time. The electrospray voltage of 2.0 kV was applied in a split-flow configuration, and spectra were collected using a top-9 data-dependent method. Survey full-scan MS spectra (m/z 400-1800) were acquired at a resolution of 70,000 with an AGC target value of 3×106 ions or a maximum ion injection time of 200 ms. The peptide fragmentation was performed via higher-energy collision dissociation with the energy set at 28 normalized collision energy (NCE). The MS/MS spectra were acquired at a resolution of 17,500, with a targeted value of 2×104 ions or maximum integration time of 200 ms. The ion selection abundance threshold was set at 8.0×102 with charge state exclusion of unassigned and z =1, or 6-8 ions and dynamic exclusion time of 30 seconds.

For database search and label-free quantitative analysis, peptide spectrum matching of MS/MS spectra of each file was searched against the human database (UniProt) using the Sequest algorithm within Proteome Discoverer v 2.3 software (Thermo Fisher Scientific, San Jose, CA). The Sequest database search was performed with the following parameters: trypsin enzyme cleavage specificity, 2 possible missed cleavages, 10 ppm mass tolerance for precursor ions, 0.02 Da mass tolerance for fragment ions. Search parameters permitted variable modification of methionine oxidation (+15.9949 Da) and static modification of carbamidomethylation (+57.0215 Da) on cysteine. Peptide assignments from the database search were filtered down to a 1% FDR. The relative label-free quantitative and comparative among the samples were performed using the Minora algorithm and the adjoining bioinformatics tools of the Proteome Discoverer 2.3 software. To select proteins that show a statistically significant change in abundance between two groups, a threshold of 1.5-fold change with p-value (0.05) were selected.

###### Cytokine Profiling

Multiplex assays were performed, as described previously in Kitajima et al., 2019, using Human Cytokine/Chemokine Magnetic Bead Panel (Cat.# HCYTMAG-60K-PX30) on a Luminex MAGPIX system (Merck Millipore). Briefly, conditioned media concentration levels (pg/ mL) of each protein were determined by 5-parameter curve fitting models. Average of two replicate fold changes relative to the corresponding control were calculated and plotted as log^2^FC.

###### Animal studies

Mouse experiments were conducted in accordance with a Kanazawa University Institutional Animal Care and Use Committee–approved protocol. Female KSN/Slc nude mice were subcutaneously injected on their armpits of right anterior limbs with 5×10^5^ cells of each group of H69AR cells, and then monitored daily for tumor growth for 2∼3 weeks until the tumor reaches to 1500mm^3^. Tumor volume were measured with a caliper and calculated by the formula, tumor size = *ab^2^/2*. *a* is the larger and *b* is the smaller of the two dimensions. When the xenografted tumors grew up to a mean tumor volume of around 250 mm^3^, tumor-bearing mice were intraperitoneally injected with various doses of Cisplatin (1mg kg^-1^) or saline every other day for a total of six times (Li et al., 2013). Tumor volume and body weight were monitored over time. Tumor progression and regression were monitored daily. The tumor-bearing mice were sacrificed 39 days after treatment, and the xenografts, lung and liver tissues were removed and subjected to RNA extraction for RT-qPCR or to 10% PFA fixation for Immunohistochemistry for hematoxylin and eosin staining.

RT-qPCR was as performed against the genes expressed in the vector such as human DDX4, NanoLuc, or Cas9 gene using primers or Taqman probe (Thermo Fisher Scientific) (primer sequences are summarized in the resources table). As controls, RNAs of the same tissue types derived from non-tumor bearing mice were used. RNA extraction was performed using the RNeasy Mini Kit (Qiagen, Cat.# 74106). RNA samples (1 μg) were reverse-transcribed using Super-Script III First-Strand Synthesis SuperMix (Thermo Fisher Scientific, Cat.# 1683483). Quantitative real-time PCR was performed using Power SYBR Green PCR Master Mix (Thermo Fisher Scientific, Cat.# 4367659). Values represent the average of three technical replicates from at least two independent experiments (biological replicates).

###### Clinical data mining

From 79 tumor samples which were previously published in Jiang et al. (2016), fastq files were fetched from GEO (accession number GSE60052) and the clinical data was downloaded from the supplementary data provided in the paper. The other tumor data (ex. AML, ESCA, LUAD) were acquired from TCGA or TARGET by using a R package called TCGAbiolinks package. 48 SCLC patient samples had survival data which among them 40 samples were treatment naïve. Non-tumor patient data were removed from all the tumor data.he samples were further divided into low and high DDX4 based on maximally ranked statistics. The survminer (version 0.4.6) and survival (3.1-8 version) R packages were used to perform survival analysis. EdgeR package in DEBrowser (version 1.2.4) was used to perform differential gene expression analysis at an adjusted p-value cut-off of <0.01 and fold change>1.5. Counts were filtered in a way that the counts distribution follows normal distribution. The EnhancedVolcano R package was used to visualize the results of differentially expressed genes. Furthermore, these expressed genes were processed to gene sets enrichment and/or functional analysis on the following gene sets (inflammatory response and immune response, germline factors, and piwiRNA biogenesis). The results of enrichment and/or functional analysis were visualized by clusterProfiler.

###### Statistical Analysis

All images were analyzed by Image-J (NIH), and statistical significance was performed by PRISM (GraphPad) using one-way or two-way ANOVA followed by Tukey post hoc test. P values less than 0.05 were considered significant. Asterisks were used to indicate significance corresponding with * is p < 0.05, ** is p < 0.01, *** is p < 0.001, **** is p < 0.0001. Columns represent means ± SD or SEM.

## REFERENCES

1. Aman A, Piotrowski T. Cell migration during morphogenesis. Dev Biol. 341(1): 20–33. 2010. PMID:19914236

2. Barretina J, Caponigro G, Stransky N, Venkatesan K, Margolin AA, Kim S, Wilson CJ, Lehár J, Kryukov GV, Sonkin D, Reddy A, Liu M, Murray L, Berger MF, Monahan JE, Morais P, Meltzer J, Korejwa A, Jané-Valbuena J, Mapa FA, Thibault J, Bric-Furlong E, Raman P, Shipway A, Engels IH, Cheng J, Yu GK, Yu J, Aspesi P Jr, de Silva M, Jagtap K, Jones MD, Wang L, Hatton C, Palescandolo E, Gupta S, Mahan S, Sougnez C, Onofrio RC, Liefeld T, MacConaill L, Winckler W, Reich M, Li N, Mesirov JP, Gabriel SB, Getz G, Ardlie K, Chan V, Myer VE, Weber BL, Porter J, Warmuth M, Finan P, Harris JL, Meyerson M, Golub TR, Morrissey MP, Sellers WR, Schlegel R, Garraway LA. The Cancer Cell Line Encyclopedia enables predictive modeling of anticancer drug sensitivity. Nature 2012. 483: 603–7. Erratum in 492(7428): 290.

3. Bartoli M, Gu X, Tsai NT, Venema RC, Brooks SE, Marrero MB, Caldwell RB. Vascular endothelial growth factor activates STAT proteins in aortic endothelial cells. J Biol Chem. 2000. 275(43):33189–92.

4. Burger JA, Stewart DJ. CXCR4 chemokine receptor antagonists: perspectives in SCLC. Expert Opin Investig Drugs 2009.18(4):481-90.

5. Bure IV, Nemtsova MV, Zaletaev DV. Roles of E-cadherin and Noncoding RNAs in the Epithelial-mesenchymal Transition and Progression in Gastric Cancer. Int J Mol Sci. 2019. 20(12). pii: E2870. DOI: 10.3390/ijms20122870. Review.

6. Calbo J, van Montfort E, Proost N, van Drunen E, Beverloo HB, Meuwissen R, Berns A. A functional role for tumor cell heterogeneity in a mouse model of small cell lung cancer. Cancer Cell. 19(2): 244–56. 2011. PMID: 21316603

7. Carrera P, Johnstone O, Nakamura A, Casanova J, J€ackle H, Lasko P. VASA mediates translation through interaction with a Drosophila yIF2 homolog. Mol Cell 5:181–7. 2000. PMID: 10678180

8. Chandrakesan P, Yao J, Qu D, May R, Weygant N, Ge Y, Ali N, Sureban SM, Gude M, Vega K, Bannerman-Menson E, Xia L, Bronze M, An G, Houchen CW. Dclk1, a tumor stem cell marker, regulates pro-survival signaling and self-renewal of intestinal tumor cells. Mol Cancer. 16(1):30. 2017. PMCID: PMC5286867

9. Davey RA, Grossmann M. Androgen Receptor Structure, Function, and Biology: From Bench to Bedside. Clin Biochem Rev. 37(1): 3–15. 2016. PMCID: PMC4810760

10. Dehghani, M., and Lasko, P. (2015). C-terminal residues specific to Vasa among DEAD-box helicases are required for its functions in piRNA biogenesis and embryonic patterning. Dev Genes Evol. 226, 401–412.

11. Doitsidou M, Reichman-Fried M, Stebler J, Köprunner M, Dörries J, Meyer D, Esguerra CV, Leung T, Raz E. Guidance of primordial germ cell migration by the chemokine SDF-1. Cell 111, 647–659. 2002. PMID:12464177

12. Dick FA, Rubin SM. Molecular mechanisms underlying RB protein function. Nat Rev Mol Cell Biol.14(5): 297– 306. 2013. PMCID:PMC4754300

13. Drapkin BJ, George J, Christensen CL, Mino-Kenudson M, Dries R, Sundaresan T, Phat S, Myers DT, Zhong J, Igo P, Hazar-Rethinam MH, LiCausi JA, Gomez-Caraballo M, Kem M, Jani KN, Azimi R, Abedpour N, Menon R, Lakis S, Heist RS, Büttner R, Haas S, Sequist LV, Shaw AT, Wong KK, Hata AN, Toner M, Maheswaran S, Haber DA, Peifer M, Dyson N, Thomas RK, Farago AF. Genomic and functional fidelity of small cell lung cancer patient-derived xenografts. Cancer Discov. 8(5):600–615. PMID: 29483136 PII: CD-17-0935. DOI: 10.1158/2159-8290. 2018.

14. Edwardson DW, Boudreau J, Mapletoft J, Lanner C, Kovala AT, Parissenti AM. Inflammatory cytokine production in tumor cells upon chemotherapy drug exposure or upon selection for drug resistance. PLoS One. 12(9):e0183662. 2017.

15. Gorelik L, Flavell RA. Immune-mediated eradication of tumors through the blockade of transforming growth factor-beta signaling in T cells. Nat Med 7:1118–22. 2001. PMID:11590434

16. Gustafson EA, Wessel GM. DEAD-box helicases: post translational regulation and function. Biochem Biophys Res Commun 395: 1–6. 2010. PMCID: PMC2863303

17. Han C, Fu J, Liu Z, Huang H, Luo L, and Yin Z: Dipyrithione inhibits IFN-gamma-induced JAK/STAT1 signaling pathway activation and IP-10/CXCL10 expression in RAW264.7 cells. Inflamm Res 59: 809–816, 2010.

18. Hanahan D, Weinberg RA. Hallmarks of cancer: the next generation. Cell 144: 646–74. 2011. PMID: 21376230

19. Hartmann TN, Burger M, Burger JA. The role of adhesion molecules and chemokine receptor CXCR4 (CD184) in small cell lung cancer. J Biol Regul Homeost Agents. 2004. 18(2):126-30.

20. Hartmann TN, Burger JA, Glodek A, Fujii N, Burger M. CXCR4 chemokine receptor and integrin signaling co-operate in mediating adhesion and chemoresistance in small cell lung cancer (SCLC) cells. Oncogene. 2005. 24(27): 4462–71.

21. Hashimoto H, Sudo T, Mikami Y, Otani M, Takano M, Tsuda H, Itamochi H, Katabuchi H, Ito M, Nishimura R. Germ cell-specific protein VASA is over-expressed in epithelial ovarian cancer and disrupts DNA damage-induced G2 checkpoint. Gynecol. Oncol. 111: 312–319. 2008. PMID:18805576

22. Hay B, Jan LY, Jan YN. A protein component of Drosophila polar granules is encoded by vasa and has extensive sequence similarity to ATP-dependent helicases. Cell 55: 577–87. 1988.

23. Heckmann L, Pock T, Tröndle I, Neuhaus N. The C-X-C signalling system in the rodent vs primate testis: impact on germ cellniche interaction. Reproduction. 2018. 155(5): R211–R219.

24. Heim MH. HCV Innate Immune Responses Viruses 1: 1073–1088. 2009. PMCID: PMC3185522

25. Hondele, M., Sachdev, R., Heinrich, S., Wang, J., Vallotton, P., Fontoura, B. M. A., & Weis, K. (2019). DEAD-box ATPases are global regulators of phase-separated organelles. Nature, 573(7772), 144–148. https://doi.org/10.1038/s41586-019-1502-y

26. Huang C, Li N, Li Z, Chang A, Chen Y, Zhao T, Li Y, Wang X, Zhang W, Wang Z, Luo L, Shi J, Yang S, Ren H, Hao J. Tumour derived Interleukin 35 promotes pancreatic ductal adenocarcinoma cell extravasation and metastasis by inducing ICAM1 expression. Nat Commun. 8:14035. 2017. PMCID: PMC5253665

27. Janice A, Mendizabal L, Llamazares S, Rossell D, Gonzalez C. Ectopic expression of germline genes drives malignant brain tumor growth in Drosophila. Science 330,1824–1827. 2010. PMID:21205669

28. Jiang L, Huang J, Higgs BW, Hu Z, Xiao Z, Yao X, Conley S, Zhong H, Liu Z, Brohawn P, Shen D, Wu S, Ge X, Jiang Y, Zhao Y, Lou Y, Morehouse C, Zhu W, Sebastian Y, Czapiga M, Oganesyan V, Fu H, Niu Y, Zhang W, Streicher K, Tice D, Zhao H, Zhu M, Xu L, Herbst R, Su X, Gu Y, Li S, Huang L, Gu J, Han B, Jallal B, Shen H, Yao Y. Genomic Landscape Survey Identifies *SRSF1* as a Key Oncodriver in Small Cell Lung Cancer. PLoS Genet. 12(4): e1005895. 2016.

29. Johnstone O, Lasko P. InteractionwitheIF5BisessentialforVasafunction during development. Development 131: 4167–4178. 2004. PMID: 15280213

30. Kardash E, Reichman-Fried M, Maître JL, Boldajipour B, Papusheva E, Messerschmidt EM, Heisenberg CP, Raz E. A role for Rho GTPases and cell-cell adhesion in single-cell motility in vivo. Nat Cell Biol. 12(1): 47–53. 2010. PMID:20010816

31. Kijima T, Maulik G, Ma PC, Tibaldi EV, Turner RE, Rollins B, Sattler M, Johnson BE and Salgia R. Cancer Res 62, 6304–6311. 2002. PMID:12414661

32. Kim KH, Kang YJ, Jo JO, Ock MS, Moon SH, Suh DS, Yoon MS, Park ES, Jeong N, Eo WK, Kim HY, Cha HJ. DDX4(DEAD-box polypeptide4) colocalizes with cancer stem cell marker CD133 in ovarian cancers. Biochem. Biophys. Res. Commun. 447: 315–322. 2014. PMID: 24727449

33. Knaut H, Werz C, Geisler R, Nusslein-Volhard C. A zebrafish homolog of the chemokine receptor Cxcr4 is a germ-cell guidance receptor. Nature 421, 279–282. 2003. PMID:12508118

34. Kunwar PS, Siekhaus DE, Lehmann R. In vivo migration: a germ cell perspective. Annu. Rev.Cell Dev. Biol. 22: 237–265. 2006. PMID:16774460

35. Lasko P. The DEAD-box helicase Vasa: evidence for a multiplicity of functions in RNA processes and developmental biology. Biochem Biophys Acta 1829:810–6. 2013. PMID:23587717

36. Lasko PF, Ashburner M. The product of the Drosophila gene vasa is very similar to eukaryotic initiation factor-4A. Nature 335: 611–7. 1988.

37. Li D, Zhang Y, Xie Y, Xiang J, Zhu Y, Yang J. Enhanced tumor suppression by adenoviral PTEN gene therapy combined with Cisplatin chemotherapy in small-cell lung cancer. Cancer Gene Ther. 20(4): 251–9. 2013. PMID:23470565

38. Lim, A. K., Lorthongpanich, C., Chew, T. G., Tan, C. W., Shue, Y. T., Balu, S., Gounko, N., Kuramochi-Miyagawa, S., Matzuk, M. M., Chuma, S., Messerschmidt, D. M., Solter, D., & Knowles, B. B. (2013). The nuage mediates retrotransposon silencing in mouse primordial ovarian follicles. Development, 140(18), 3819–3825. https://doi.org/10.1242/dev.099184

39. Linder P. Dead-box proteins: a family affair active and passive players in RNP-remodeling. Nucleic Acids Res 34: 4168–80. 2006.

40. Linder P, Lasko PF, Ashburner M, Leroy P, Nielsen PJ, Nishi K, Schnier J, Slonimski PP. Birth of the D-E-A-D box. Nature 337,121–122. 1989.

41. Liu N, Han H, Lasko P. Vasa promotes Drosophila germline stem cell differentiation by activating mei-P26 translation by directly interacting with a (U)-rich motif in its 3’ UTR. Genes Dev 23: 2742–52. 2010. PMID:19952109.

42. Molyneaux, KA, Stallock J, Schaible K, Wylie C. Time-lapse analysis of living mouse germ cell migration. Dev. Biol. 240, 488–498. 2001.

43. Moreira-Pais A, Ferreira R, Gil da Costa R. Platinum-induced muscle wasting in cancer chemotherapy: Mechanisms and potential targets for therapeutic intervention. Life Sci. 208:1–9. Review.

44. Nagasawa T. CXC chemokine ligand 12 (CXCL12) and its receptor CXCR4. J Mol Med (Berl). 2014. 92(5):433–9.

45. Nakad R, Schumacher B. DNA Damage Response and Immune Defense: Links and Mechanisms. Front Genet. 7:147. 2016. PMCID: PMC4977279

46. Nakamura Y, Yamamoto Y, Usui F, Mushika T, Ono T, Setioko AR, Takeda K, Nirasawa K, Kagami H, Tagami T. Migration and proliferation of primordial germ cells in the early chicken embryo. Poult Sci. 86(10):2182–93. 2007.

47. Nangia-Makker P, Hogan V, Raz A. Galectin-3 and cancer stemness. Glycobiology. 2018. 28(4):172–181. DOI: 10.1093/glycob/cwy001.

48. Ni L, Lu J. Interferon-gamma in cancer immunotherapy. Cancer Med. 7(9):4509–4516. 2018.

49. Nott, T. J., Petsalaki, E., Farber, P., Jervis, D., Fussner, E., Plochowietz, A., Craggs, T. D., Bazett-Jones, D. P., Pawson, T., Forman-Kay, J. D., & Baldwin, A. J. (2015). Phase transition of a disordered nuage protein generates environmentally responsive membraneless organelles. Mol Cell, 57(5), 936–947. https://doi.org/10.1016/j.molcel.2015.01.013

50. Pestova TV, Lomakin IB, Lee JH, Choi SK, Dever TE, Hellen CUT. The joining of ribosomal subunits in eukaryotes requires eIF5B. Nature 403, 332–335. 2000.

51. Poon J, Wessel GM, Yajima M. An unregulated regulator: ectopic Vasa expression and tumorigenesis. Develop Biol 415: 24–32. 2016. PMCID:PMC4902722

52. Raz E. The function and regulation of vasa-like genes in germ-cell development. Genome Biol 1: 1017.1. 2000.

53. Richardson BE, Lehmann R. Mechanisms guiding primordial germ cell migration: strategies from different organisms. Nat Rev Mol Cell Biol 11: 37–49. 2010.

54. Riedl J, Crevenna AH, Kessenbrock K, Yu JH, Neukirchen D, Bista M, Bradke F, Jenne D, Holak TA, Werb Z, Sixt M, Wedlich-Soldner R. Lifeact: a versatile marker to visualize F-actin. Nat Methods. 5(7): 605–7. 2008. PMCID:PMC2814344

55. Roth S, Rottach A, Lotz-Havla AS, Laux V, Muschaweckh A, Gersting SW, Muntau AC, Hopfner KP, Jin L, Vanness K, Petrini JH, Drexler I, Leonhardt H, Ruland J. 2014. Rad50 CARD9 interactions link cytosolic DNA sensing to IL-1β production. Nat Immunol. 15(6):538–45.

56. Rothenberger NJ, Somasundaram A, Stabile LP. The Role of the Estrogen Pathway in the Tumor Microenvironment. Int J Mol Sci. 19(2). 2018. PMCID:PMC5855833

57. Sanjana NE, Shalem O, Zhang F. Improved lentiviral vectors and genome-wide libraries for CRISPR screening. Nature Methods 11(8): 783–784. 2014.

58. Sen T, Rodriguez BL, Chen L, Corte CMD, Morikawa N, Fujimoto J, Cristea S, Nguyen T, Diao L, Li L, Fan Y, Yang Y, Wang J, Glisson BS, Wistuba II, Sage J, Heymach JV, Gibbons DL, Byers LA. Targeting DNA Damage Response Promotes Antitumor Immunity through STING-Mediated T-cell Activation in Small Cell Lung Cancer. Cancer Discov. 9(5):646–661. 2019.

59. Schudrowitz, N., Takagi, S., Wessel, G.M., Yajima, M. Germline factor DDX4 functions in blood-derived cancer cell phenotypes. Cancer Sci.108: 1612–1619. 2017. PMCID: PMC5543511

60. Schwager EE, Meng Y, Extavour CG. Vasa and piwi are required for mitotic integrity in early embryogenesis in the spider Parasteatodate pidariorum. Dev. Biol. 402(2): 276–290. 2014. PMID:25257304

61. Sengoku T, Nureki O, Nakamura A, Kobayashi S, Yokoyama S. Structural basis for RNA unwinding by the DEAD-box protein Drosophila Vasa. Cell 125: 287–300. 2006.

62. Sharma A, Yeow WS, Ertel A, Coleman I, Clegg N, Thangavel C, Morrissey C, Zhang X, Comstock CE, Witkiewicz AK, Gomella L, Knudsen ES, Nelson PS, Knudsen KE. The retinoblastoma tumor suppressor controls androgen signaling and human prostate cancer progression. J Clin Invest. 120(12): 4478–92. 2010. PMCID: PMC2993601

63. Shimaoka K, Mukumoto Y, Tanigawa Y, Komiya T. Xenopus Vasa Homolog XVLG1 is Essential for Migration and Survival of Primordial Germ Cells. Zoolog Sci. 34(2):93–104. 2017. PMID:28397605

64. Simpson AJ, Caballero OL, Jungbluth A, Chen YT, Old LJ. Cancer/testis antigens, gametogenesis, and cancer. Nat Rev Cancer 5: 615–625. 2005.

65. Swaim CD, Scott AF, Canadeo LA, Huibregtse JM. Extracellular ISG15 Signals Cytokine Secretion Through the LFA-1 Integrin Receptor. Mol Cell 68(3): 581–590.e5. 2017.

66. Tantin D, Gemberling M, Callister C, Fairbrother WG. High-throughput biochemical analysis of in vivo location data reveals novel distinct classes of POU5F1(Oct4)/DNA complexes. Genome Res.18(4): 631–9. 2008. PMCID: PMC2279250

67. Trenti A, Tedesco S, Boscaro C, Trevisi L, Bolego C, Cignarella A. Estrogen, Angiogenesis, Immunity and Cell Metabolism: Solving the Puzzle. Int J Mol Sci.19(3). 2018. PMCID:PMC5877720

68. Wagner DE, Ho JJ, Reddien PW. Genetic regulators of a pluripotent adult stem cell system in planarians identified by RNAi and clonal analysis. Cell Stem Cell 10: 299–311. 2012. PMCID:PMC3338251

69. Wang D, Kennedy S, Conte DJ, Kim JK, Gabel HW, Kamath RS, Mello CC, Ruvkun G. Somatic misexpression of germline P granules and enhanced RNA interference in retinoblastoma pathway mutants. Nature 436: 593–597. 2005.

70. Yachida S, Iacobuzio-Donahue CA. Evolution and dynamics of pancreatic cancer progression. Oncogene 32: 5253–60. 2013. PMCID:PMC3823715

71. Yajima M, Umeda R, Fuchikami T, Kataoka M, Sakamoto N, Yamamoto T, Akasaka K. Implication of HpEts in gene regulatory networks responsible for specification of sea urchin skeletogenic primary mesenchyme cells. Zool. Sci. 27:638–646. 2010.

72. Yajima M, Wessel GM. The DEAD-box RNA helicase Vasa functions in embryonic mitotic progression in the sea urchin. Development 138: 2217–22. 2011. PMCID: PMC3091493

73. Yajima M, Fairbrother W, Wessel GM. ISWI contributes to ArsI insulator function in development of the sea urchin. Development 139: 3613–3622. 2012. PMCID:PMC3436113

74. Yajima M, Wessel GM. The germline factor Vasa functions broadly in somatic cells: mRNA clustering, translational regulation, and wound healing. Development 142(11),1960–1970. 2015. PMCID:PMC4460737

75. Zhang P, Hu P, Shen H, Yu J, Liu Q, Du J. Prognostic role of Twist or Snail in various carcinomas: a systematic review and meta-analysis. Eur J Clin Invest. 2014. 44(11):1072–94. doi:10.1111/eci.12343. Review.

76. Zhang Z, Zhou Y, Qian H, Shao G, Lu X, Chen Q, Sun X, Chen D, Yin R, Zhu H, Shao Q, Xu W. Stemness and inducing differentiation of small cell lung cancer NCI-H446 cells. Cell Death Dis 4:e633. 2013. PMCID: PMC3674360.

77. Zheng L, Meng Y, Campbell JL, Shen B. Multiple roles of DNA2 nuclease/helicase in DNA metabolism, genome stability, and human diseases. Nucleic Acids Res. 48(1): 16–35. 2020.

78. Zhou X, Wei J, Chen F, Xiao X, Huang T, He Q, Wang S, Du C, Mo Y, Lin L, Xie Y, Wei L, Lan Y, Murata M, Huang G, Ernberg I, Matskova L, Zhang Z. Epigenetic downregulation of the ISG15–conjugating enzyme UbcH8 impairs lipolysis and correlates with poor prognosis in nasopharyngeal carcinoma. Oncotarget. 6(38): 41077–41091. 2015.

## References for Supplementary Materials

79. Ahsan N, Belmont J, Chen Z, Clifton JG, Salomon AR. Highly reproducible improved label-free quantitative analysis of cellular phosphoproteome by optimization of LC-MS/MS gradient and analytical column construction. J Proteomics 2017, 165: 69–74.

80. Chen D, Cao Y, Li H, Kim D, Ahsan N, Thelen J, Stacey G. Extracellular ATP elicits DORN1-mediated RBOHD phosphorylation to regulate stomatal aperture. Nat Commun. 8(1): 2265. 2017.

81. Kitajima S, Asahina H, Chen T, Guo S, Quiceno LG, Cavanaugh JD, Merlino AA, Tange S, Terai H, Kim JW, Wang X, Zhou S, Xu M, Wang S, Zhu Z, Thai TC, Takahashi C, Wang Y, Neve R, Stinson S, Tamayo P, Watanabe H, Kirschmeier PT, Wong KK, Barbie DA. Overcoming Resistance to Dual Innate Immune and MEK Inhibition Downstream of KRAS. Cancer Cell 34(3):439-452. 2018.

82. Kitajima S, Ivanova E, Guo S, Yoshida R, Campisi M, Sundararaman SK, Tange S, Mitsuishi Y, Thai TC, Masuda S, Piel BP, Sholl LM, Kirschmeier PT, Paweletz CP, Watanabe H, Yajima M, Barbie DA. Suppression of STING Associated with LKB1 Loss in KRAS -Driven Lung Cancer. Cancer Discov. 9:34–45. 2019.

83. Shalem O, Sanjana NE, Hartenian E, Shi X, Scott DA, Mikkelsen T, Heckl D, Ebert BL, Root DE, Doench JG, Zhang F. Genome-scale CRISPR-Cas9 knockout screening in human cells. Science 343: 83–87. 2014.

84. Schudrowitz, N., Takagi, S., Wessel, G.M., Yajima, M. Germline factor DDX4 functions in blood-derived cancer cell phenotypes. Cancer Sci.108: 1612–1619. 2017.

85. Yu K, Sabelli A, DeKeukelaere L, Park R, Sindi S, Gatsonis CA, Salomon A. Integrated platform for manual and high-throuput statistical validation of tadem mass spectra. Proteomics 9(11): 3115–25. 2009.

86. Yu K, Salomon A. PeptideDepot: Flexible relational database for visual analysis of quantitative proteomic data and integration of existing protein information. Proteomics 9(23): 5350–58. 2009.

87. Yu K, Salomon A. HTAPP: High-throughput autonomous proteomic pipeline. Proteomics 10, 2113–2122. s2010.

88. Jiang, Liyan, et al. “Genomic landscape survey identifies SRSF1 as a key oncodriver in small cell lung cancer.” PLoS genetics 12.4 (2016).

89. Kucukural, A., Yukselen, O., Ozata, D.M. et al. DEBrowser: interactive differential expression analysis and visualization tool for count data. BMC Genomics 20, 6 (2019). https://doi.org/10.1186/s12864-018-5362-x

90. Kerley-Hamilton, J., Pike, A., Li, N. et al. A p53-dominant transcriptional response to cisplatin in testicular germ cell tumor-derived human embyronal carcinoma. Oncogene 24, 6090–6100 (2005). https://doi.org/10.1038/sj.onc.1208755

91. Pillai, R.S. and Chuma, S. (2012), piRNAs and their involvement in male germline development in mice. Development, Growth & Differentiation, 54: 78–92. https://doi.org/10.1111/j.1440-169X.2011.01320.x

92. Siomi MC, Sato K, Pezic D, Aravin AA. PIWI-interacting small RNAs: the vanguard of genome defence. Nat Rev Mol Cell Biol. 2011 Apr;12(4):246–58. doi: 10.1038/nrm3089. PMID: 21427766.

93. Mark A. Lindsay, Sam Griffiths-Jones, Kaoru Sato, Mikiko C. Siomi; Piwi-interacting RNAs: biological functions and biogenesis. Essays Biochem 3 May 2013; 54 39–52. doi: https://doi.org/10.1042/bse0540039

